# Application of an Optimized Annotation Pipeline to the *Cryptococcus Deuterogattii* Genome Reveals Dynamic Primary Metabolic Gene Clusters and Genomic Impact of RNAi Loss

**DOI:** 10.1101/2020.09.01.278374

**Authors:** Patrícia Aline Gröhs Ferrareze, Corinne Maufrais, Rodrigo Silva Araujo Streit, Shelby J. Priest, Christina Cuomo, Joseph Heitman, Charley Christian Staats, Guilhem Janbon

**Author notes:** Corresponding author Address of the corresponding author: Guilhem Janbon, Institut Pasteur, Unité Biologie des ARN des Pathogènes Fongiques, Département de Mycologie, 25 rue du Dr Roux,75015, Paris, France. Both authors should be considered as senior authors (Charley Christian Staats, Guilhem Janbon).

## Abstract

Evaluating the quality of a *de novo* annotation of a complex fungal genome based on RNA-seq data remains a challenge. In this study, we sequentially optimized a Cufflinks-CodingQuary based bioinformatics pipeline fed with RNA-seq data using the manually annotated model pathogenic yeasts *Cryptococcus neoformans* and *Cryptococcus deneoformans* as test cases. Our results demonstrate that the quality of the annotation is sensitive to the quantity of RNA-seq data used and that the best quality is obtained with 5 to 10 million reads per RNA-seq replicate. We also demonstrated that the number of introns predicted is an excellent *a priori* indicator of the quality of the final *de novo* annotation. We then used this pipeline to annotate the genome of the RNAi-deficient species *Cryptococcus deuterogattii* strain R265 using RNA-seq data. Dynamic transcriptome analysis revealed that intron retention is more prominent in *C. deuterogattii* than in the other RNAi-proficient species *C. neoformans* and *C. deneoformans*. In contrast, we observed that antisense transcription was not higher in *C. deuterogattii* than in the two other *Cryptococcus* species. Comparative gene content analysis identified 21 clusters enriched in transcription factors and transporters that have been lost. Interestingly, analysis of the subtelomeric regions in these three annotated species identified a similar gene enrichment, reminiscent of the structure of primary metabolic clusters. Our data suggest that there is active exchange between subtelomeric regions, and that other chromosomal regions might participate in adaptive diversification of *Cryptococcus* metabolite assimilation potential.

## Introduction

In recent years, we have seen an astonishing multiplication of fungal genome sequences (James *et al.* 2020). Long-read sequencing and adapted bioinformatics tools are quickly improving as well. It is expected that telomere-to-telomere whole-genome sequencing will soon become standard for reference genomes of diverse organisms (Giordano *et al.* 2017; Dal Molin *et al.* 2018; Yadav *et al.* 2018). Yet, fungal genomes remain difficult to annotate. Historically, most annotation tools have relied upon comparative genomics, but other pipelines utilize RNA-seq data or a combination of both approaches to propose gene annotation models (Cantarel *et al.* 2008; Haas *et al.* 2011; Min *et al.* 2017; Haridas *et al.* 2018). These pipelines are very efficient in intron-poor species, at least for predicting coding regions. For instance, a recent MAKER-based optimized pipeline tested on 39 budding yeast genomes missed only 3.9% of genes and 4.8% of exons, on average (Shen *et al.* 2018). However, the results were poorer in intron-rich species, for which gene annotation is challenging. Even when RNA-seq data are available, it is still very difficult to correctly predict the exon-intron structure primarily because fungal exons can be extremely short (Janbon *et al.* 2014), but also because these genomes are compact. Thus, when tested on fungal data sets, *de novo* transcriptome assemblers like Trinity (Grabherr *et al.* 2011) or Cufflinks (Trapnell *et al.* 2010) tend to predict very large transcripts with no biological relevance. Nevertheless, several pipelines have been published and sequencing centers like the Joint Genome Institute (JGI) and the Broad Institute have developed specialized pipelines to produce annotation drafts, which are very useful in large-scale comparison analyses (Haas *et al.* 2011; Haridas *et al.* 2018).

Some methods, like the construction of large deletion collections, or precise analysis of gene content needs more precise annotation, and the annotation strategy applied will depend on the goal of the research (Mudge and Harrow 2016). Manual curation of a pre-annotated genome will likely result in the highest-quality gene prediction. Some tools, like Artemis (Carver *et al.* 2012) and Apollo (Dunn *et al.* 2019), have been used to manually curate annotation, but they are time consuming even when several annotators are implemented. Without manual curation, it is impossible to anticipate the results from an annotation bioinformatics pipeline fed with RNA-seq data. Typically, the quality of the prediction will depend on the diversity, quantity, and quality of the data, but no *a priori* indicator exists to determine if the *de novo* gene prediction is accurate.

Pathogenic *Cryptococcus* species are basidiomycete yeasts, which cause nearly 200,000 deaths annually around the world (Kwon-Chung *et al.* 2014). There are currently eight recognized pathogenic species of *Cryptococcus* (Hagen *et al.* 2015; Farrer et al., 2019). Manual annotation of the *Cryptococcus neoformans* and *Cryptococcus deneoformans* reference genomes revealed complex and dynamic transcriptomes (Janbon *et al.* 2014; Wallace *et al.* 2020). These annotations were recently completed through precise identification of the transcript leader (TL) and 3’UTR sequences through TSS-seq and 3UTR-seq analyses; these annotations are likely the most complete and detailed annotations in intron-rich fungi (Wallace *et al.* 2020). With 99.5% of 6,795 annotated coding genes containing introns, five to six introns per coding gene, and 37,832 introns in total, an automatic annotation of these genomes would be considered highly challenging even with the large sets of RNA-seq data that have been produced (Wilm *et al.* 2007; Janbon 2018; Wallace *et al.* 2020).

In this study, we compared the performances of three annotation pipelines fed with RNA-seq data. We gradually optimized the quality of the *de novo* annotation using the well-annotated genomes of *C. neoformans* and *C. deneoformans* as ground-truth inputs. We found that the quantity of data used should not be too large and that the number of introns predicted had a positive, linear relationship with the quality of the *de novo* annotation. We used this pipeline to re-annotate the reference genome of the RNAi-deficient *Cryptococcus deuterogattii* strain R265 using RNA-seq data. Analysis of the transcriptome dynamics of these three *Cryptococcus* species revealed that although the sense/antisense transcript ratio is similar across all three species, intron retention is higher in *C. deuterogattii*. Comparative gene content analysis identified a list of genes that are absent or largely truncated in R265, many of which have been implicated in RNAi-mediated silencing in *Cryptococcus* species. Finally, we also identified several primary metabolic gene clusters (MGCs) that are absent in R265 and associated this loss with the subtelomeric gene content. Our data suggest an active exchange of MGCs between subtelomeric regions and more central regions of the genome. This exchange might contribute to the adaptive diversification of metabolite assimilation potential in *Cryptococcus*.

## MATERIALS AND METHODS

### RNA-Seq sample and data production

RNA-seq libraries from four growth conditions (exponential phase at 30°C, + exponential phase at 37°C, stationary phase at 30°C, and stationary phase at 37°), conducted in triplicate, of *C. neoformans* H99 and *C. deneoformans* JEC21 used in this study have been previously described (Wallace et al. 2020). The *C. deuterogattii* R265 strain was grown in YPD at 30°C and 37°C under agitation to exponential or early stationary phase as previously described (Wallace et al. 2020). Briefly, early stationary phase was obtained after 18 h of growth (final OD 600= 15) starting from at OD600 = 0.5. Each *Cryptococcus* cell preparation was spiked in with one tenth (OD/OD) of *S. cerevisiae* strain FY834 cells grown in YPD at 30°C in stationary phase. Cells were washed, snap frozen and used to prepare RNA and total DNA samples. Biological triplicates were prepared in each condition. For RNA-seq, strand-specific, paired-end cDNA libraries were prepared from 10 μg of total RNA following poly-A purification using the TruSeq Stranded mRNA kit (Illumina) according to manufacturer’s instructions. cDNA fragments of ~400 bp were purified from each library and confirmed for quality by Bioanalyzer (Agilent). DNA-Seq libraries were prepared using the kit TruSeq DNA PCR-Free (Illumina). Then, 100 bases were sequenced from both ends using an Illumina HiSeq2500 instrument according to the manufacturer’s instructions (Illumina). For the mating condition, total RNA was isolated (in biological triplicates) from a *C. neoformans* cross between the congenic mating partners H99 (*MAT*α) and YL99 (*MAT***a**) (Semighini et al., 2011) or a *C. deuterogattii* cross between the congenic mating partners R265 (*MAT*α) and AIR265 (*MAT***a**) (Zhu et al. 2013). Briefly, overnight cultures were grown under standard laboratory conditions in YPD at 30°C. Overnight cultures were diluted to an OD600 = 1.0, and cells from both strains were mixed, spotted onto V8 (pH = 5) mating medium, and incubated in the dark at room temperature for 48 h. Cells were scraped from mating plates, snap frozen, and RNA was isolated using Trizol following the manufacturer’s protocol. RNA quality was confirmed by Bioanalyzer (Agilent) and RNA samples were depleted of ribosomal RNA with the Ribo-Zero Gold rRNA Removal Kit for Yeast (Illumina). Strand-specific, paired-end cDNA libraries were prepared using the TruSeq Stranded mRNA kit (Illumina), and 150 bases were sequenced from both ends using an Illumina HiSeq4000 instrument according to the manufacturer’s instructions (Illumina).

### RNA-Seq library trimming and rRNA cleaning

The paired reads from the RNA-seq libraries were trimmed for low quality reads and Illumina TruSeq adapters were removed with Cutadapt v1.9.1 (Martin 2011) with the following parameters: --trim-qualities 30 –e (maximum error rate) 0.1 --times 3 --overlap 6 --minimum-length 30. The cleaning of rRNA sequences was performed with Bowtie2 v2.3.3 (Langmead and Salzberg 2012) with default parameters; unmapped paired reads were reported using option --un-conc to identify reads that did not align with rRNA sequences.

### RNA-Seq library mapping

The cleaned reads from RNA-seq paired-end libraries from *C. neoformans* H99, *C. deneoformans* JEC21, and *C. deuterogattii* R265 were mapped against their reference genomes (NCBI Genome Assemblies GCA_000149245.3, GCA_000091045.1 and GCA_002954075.1) with Tophat2 v2.0.14 (Kim *et al.* 2013) and the following parameters: minimum intron length 30; minimum intron coverage 30; minimum intron segment 30; maximum intron length 4000; maximum multihits 1; microexon search; and library-type fr-firststrand or fr-secondstrand (according to the RNA-seq library).

### Pipeline selection

The RNA-seq mapped reads from *C. neoformans* H99 and *C. deneoformans* JEC21 from the EXPO30 condition (exponential growth at 30 C) were tested in the three pipelines for gene prediction. BRAKER1 (Hoff *et al.* 2016) was performed with the default parameters plus the exclusion of alternative transcripts (--alternatives-from-evidence=false) using the three replicates (A, B, and C) as RNA-seq source. Cuff-CQ (Cufflinks v2.1.1 ((Trapnell *et al.* 2010)) /Coding Quarry v2.0 (Testa *et al.* 2015)) and C3Q (Cufflinks v2.1.1/Cuffmerge/Coding Quarry v2.0) were tested with the basic parameters: minimum intron length (30); maximum intron length (4000); minimum isoform fraction (0.9); and overlap radius (10). The merged BAM file generated by the three replicates (A, B, and C) and used in the Cuff-CQ pipeline was obtained with Samtools *merge.* C3Q was performed separately for the three BAM files; the GTF files generated by the three predictions (for replicates A, B, and C) were then combined by Cuffmerge and the resulting transcripts were processed by CodingQuarry. The evaluation of the pipeline sensitivity and precision for gene prediction was performed by comparing the predicted annotations against the H99 and JEC21 reference annotations (Wallace et al., 2020) with the GFFCompare program (Pertea and Pertea 2020).

*For a better understanding, the C3Q pipeline with the basic Cufflinks parameters is named as C3Q1 protocol in the results section.*

### Cufflinks parameters selection

The selection of the best Cufflinks parameter combination was also performed with EXPO30 RNA-seq libraries from *C. neoformans* H99 and *C. deneoformans* JEC21 according to the C3Q pipeline. For this, the Cufflinks transcript assembly generated for each EXPO30 replicate (A, B, and C) was tested with fixed and variable parameter combinations (Table 1). Subsequently, as established for the C3Q pipeline, the predicted GTFs were merged and processed by CodingQuarry. All combinations include minimum intron length 30; maximum intron length 4000; and minimum isoform fraction 0.9; since we want to remove all isoforms. The variable parameters include: pre-mRNA fraction 0.15 to 1.0; small anchor fraction 0.0; minimum fragments per transfag 1; overlap radius 1, 10 and 100; 3’ trimming (--trim-3-avgcov-thresh and –trim-3-dropoff-frac) 0. The evaluation of the Cufflinks parameters for sensitivity and specificity for gene prediction was performed by comparison of the predicted annotations against the H99 and JEC21 reference annotations with the GFFCompare program.

**Table 1.**

|  | **Cufflinks parameter combinations** |
| --- | --- |
| **A** | --max-intron-length 4000 --min-intron-length 30 --min-isoform-fraction 0.9 |
| **B** | --max-intron-length 4000 --min-intron-length 30 --min-isoform-fraction 0.9 --min-frags-per-transfag 1 |
| **C** | --max-intron-length 4000 --min-intron-length 30 --min-isoform-fraction 0.9 --pre-mrna-fraction 0.25 |
| **D** | --max-intron-length 4000 --min-intron-length 30 --min-isoform-fraction 0.9 --overlap-radius 10 |
| **E** | --max-intron-length 4000 --min-intron-length 30 --min-isoform-fraction 0.9 --overlap-radius 100 |
| **F** | --max-intron-length 4000 --min-intron-length 30 --min-isoform-fraction 0.9 --trim-3-avgcov-thres 0 --trim-3-dropoff-frac 0.0 |
| **G** | --max-intron-length 4000 --min-intron-length 30 --min-isoform-fraction 0.9 --pre-mrna-fraction 0.25 --overlap-radius 10 --trim-3-avgcov-thresh 0 --trim-3-dropoff-frac 0.0 |
| **H** | --max-intron-length 4000 --min-intron-length 30 --min-isoform-fraction 0.9 --pre-mrna-fraction 0.25 --overlap-radius 10 --trim-3-avgcov-thresh 0 --trim-3-dropoff-frac 0.0 --min-frags-per-transfag 1 |
| **I** | --max-intron-length 4000 --min-intron-length 30 --min-isoform-fraction 0.9 --pre-mrna-fraction 0.25 --overlap-radius 1 --trim-3-avgcov-thresh 0 --trim-3-dropoff-frac 0.0 |
| **J** | --max-intron-length 4000 --min-intron-length 30 --min-isoform-fraction 0.9 --pre-mrna-fraction 0.25 --overlap-radius 10 --trim-3-avgcov-thresh 0 --trim-3-dropoff-frac 0.0 --min-frags-per-transfag 1 |
| **K** | --max-intron-length 4000 --min-intron-length 30 --min-isoform-fraction 0.9 --pre-mrna-fraction 0.25 --overlap-radius 1 --trim-3-avgcov-thresh 0 --trim-3-dropoff-frac 0.0 --small-anchor-fraction 0.0 |
| **L** | --max-intron-length 4000 --min-intron-length 30 --min-isoform-fraction 0.9 --pre-mrna-fraction 0.50 --overlap-radius 1 --trim-3-avgcov-thresh 0 --trim-3-dropoff-frac 0.0 --small-anchor-fraction 0.0 |
| **M** | --max-intron-length 4000 --min-intron-length 30 --min-isoform-fraction 0.9 --pre-mrna-fraction 0.75 --overlap-radius 1 --trim-3-avgcov-thresh 0 --trim-3-dropoff-frac 0.0 --small-anchor-fraction 0.0 |
| **N** | --max-intron-length 4000 --min-intron-length 30 --min-isoform-fraction 0.9 --pre-mrna-fraction 0.80 --overlap-radius 1 --trim-3-avgcov-thresh 0 --trim-3-dropoff-frac 0.0 --small-anchor-fraction 0.0 |
| **O** | --max-intron-length 4000 --min-intron-length 30 --min-isoform-fraction 0.9 --pre-mrna-fraction 0.90 --overlap-radius 1 --trim-3-avgcov-thresh 0 --trim-3-dropoff-frac 0.0 --small-anchor-fraction 0.0 |
| **P** | --max-intron-length 4000 --min-intron-length 30 --min-isoform-fraction 0.9 --pre-mrna-fraction 1.0 --overlap-radius 1 --trim-3-avgcov-thresh 0 --trim-3-dropoff-frac 0.0 --small-anchor-fraction 0.0 |
| **Q** | --max-intron-length 4000 --min-intron-length 30 --min-isoform-fraction 0.9 --pre-mrna-fraction 0.85 --overlap-radius 1 --trim-3-avgcov-thresh 0 --trim-3-dropoff-frac 0.0 --small-anchor-fraction 0.0 |

*For a better understanding, the C3Q pipeline with the “Q” Cufflinks parameters (selected combination) is named as C3Q2 protocol in the results section.*

### Gene predictions with H99 and JEC21 RNA-seq libraries

The validation of this gene prediction system was evaluated by applying the C3Q pipeline with the best selected Cufflinks parameters (“Q” combination) to all RNA-seq libraries from *C. neoformans* H99 and *C. deneoformans* JEC21. For H99, the fifteen libraries obtained from the five growth conditions were used (Exponential phase at 30°C, Exponential phase at 37°C; Stationary phase at 30°C, Stationary phase at 37°C and Mating). For JEC21, we tested twelve libraries obtained from four growth conditions (Exponential phase at 30°C, Exponential phase at 37°C; Stationary phase at 30°C and Stationary phase at 37°C).

The evaluation of the sensitivity and specificity for gene prediction was performed by comparison of the predicted annotations against H99 and JEC21 reference annotations with the GFFCompare program.

*For a better understanding, the C3Q pipeline with the “Q” Cufflinks parameters and the RNA-seq libraries for all the sequenced conditions (“ES3037M” for C. neoformans H99 and “ES3037” for C. deneoformans JEC21) is named as C3Q3 protocol in the results section.*

*ES3037M: Exponential phase at 30°C (EXPO30) + Exponential phase at 37°C (EXPO30) + Stationary phase at 30°C (STAT30) + Stationary phase at 37°C (STAT37) + Mating

*ES3037: Exponential phase at 30°C (EXPO30) + Exponential phase at 37°C (EXPO30) + Stationary phase at 30°C (STAT30) + Stationary phase at 37°C (STAT37)

### Effect of different conditions on predictions

To evaluate the effect of the growth conditions on the predicted annotation, we used combinations of RNA-seq libraries derived from two, three, and four of the growth conditions for *C. neoformans* H99 and of two and three of the growth conditions for *C. deneoformans* JEC21. The predictions for each combination were performed according to the C3Q pipeline and “Q” Cufflinks parameters. The evaluation of the sensitivity and specificity for gene prediction was performed by comparison of the predicted annotations against H99 and JEC21 reference annotations with the GFFCompare program.

### Evaluation of the effect of the sequencing depth on gene prediction quality

The evaluation of the effect of the sequencing depth on gene prediction was performed by down sampling the three RNA-seq libraries from the EXPO30 condition (replicates A, B, and C) with the tool *PositionBasedDownsampleSam* from Picard package (https://broadinstitute.github.io/picard/). In this analysis, *C. neoformans* H99 and *C. deneoformans* JEC21 were used. According to a random algorithm that downsamples BAM files, we used defined fractions of 1, 5, 7.5, 10, 15, 20, 30 and 40 million reads for each replicate. Subsequently, the predictions were performed according to the C3Q pipeline with the Cufflinks “Q” parameter combination using the downsampled files. Evaluation of the sensitivity and specificity of gene prediction was performed by comparison of the predicted annotations against H99 and JEC21 reference annotations with the GFFCompare program.

### Gene predictions with downsampled H99 and JEC21 RNA-seq libraries

Gene prediction using the downsampled BAM files from the RNA-seq conditions was performed according to the C3Q pipeline with “Q” Cufflinks parameters and the downsampled RNA-Seq alignment files for all *C. neoformans* H99 (Exponential phase at 30°C, Exponential phase at 37°C, Stationary phase at 30°C, Stationary phase at 37°C and Mating) and *C. deneoformans* JEC21 (Exponential phase at 30°C, Exponential phase at 37°C; Stationary phase at 30°C and Stationary phase at 37°C) growth conditions. The downsampling of each replicate to 7.5 million reads was performed with the Picard package, as previously described. Evaluation of the sensitivity and specificity of gene prediction was performed by comparison of the predicted annotations against H99 and JEC21 reference annotations with the GFFCompare program.

*For a better understanding, the C3Q pipeline with the “Q” Cufflinks parameters and the subsampled BAM files from RNA-seq libraries for all the growth conditions (“ES3037M” for C. neoformans H99 and “ES3037” for C. deneoformans JEC21) is named as C3Q4 protocol in the results section.*

### Characterization of novel and missed loci

The identification of novel and missed loci was performed with the GFFCompare program using the reference annotations from *C. neoformans* H99 and *C. deneoformans* JEC21 and the predicted C3Q gene annotations. Evaluation of the functional annotation (function, presence of domain signatures) of these sequences was performed by Blastp and Interproscan search from Blast2GO (Conesa *et al.* 2005). The expression quantification of these sequences was performed with HTSeq-count (Anders *et al.* 2014) with the following parameters *--stranded yes -f bam -r pos -t CDS*.

### Deletion of dubious novel loci from predictions

Deletion of dubious novel sequences was tested with predicted transcripts of 100 nt, 150 nt, 200 nt and 300 nt, as well as intronless sequences of 300 nt from *C. neoformans* H99 and *C. deneoformans* JEC21 C3Q predictions. The sequence deletion and evaluation of the results was performed with an in-house AWK script and the GFFCompare program. Deletion of genome-predicted sequences without supporting reads and those with low FPKM values were performed and evaluated with an in-house AWK script combined with the HTSeq-count and the GFFCompare program. Deletion of alternative transcripts from multi-transcript loci was also performed with an in-house AWK script and GFFCompare. In this process, we selected for transcripts predicted by Cufflinks with supporting RNA-seq evidence. Of these selected transcripts, the longest transcript was chosen. For the other genes predicted only from genome sequencing (without RNA-seq evidence), the longest transcript was selected.

We assessed the sensitivity and specificity of the C3Q predictions for *C. neoformans* H99 and *C. deneoformans* JEC21 against their reference annotations to analyze the effect of dubious sequence deletion. Filter combinations with the low numbers of remnant novel transcripts and smaller reduction in the prediction quality parameters were favored.

*For a better understanding, the C3Q pipeline with the “Q” Cufflinks parameters, the subsampled BAM from RNA-seq libraries for all the sequenced conditions (“ES3037M” for C. neoformans H99 and “ES3037” for C. deneoformans JEC21), and the sequence filtering (sequences up to 150 nt, intronless sequences up to 300 nt, genome-predicted sequences without reads and alternative transcripts) is named as C3Q5 protocol in the results section.*

### Retrieval of deleted and non-predicted loci

The mapping of *C. neoformans* H99 protein sequences in the *C. deneoformans* JEC21 genome and JEC21 protein sequences in the H99 genome by Exonerate v2.2.0 program (https://www.ebi.ac.uk/about/vertebrate-genomics/software/exonerate) with the following parameters (*protein2genome --percent 30 -- bestn 1 --minintron 30 – maxintron 4000 -- showalignment false --showvulgar false --showtargetgff true --refine region --subopt false)* was performed to recover sequences deleted in the previous filtering step with conserved orthology in *Cryptococcus*. For this purpose, the mapped gene coordinates matching previously predicted sequences (GFFCompare program) were used to add these deleted genes to the annotation with an in-house AWK script. The addition of non-predicted genes was performed by comparing the mapped protein sequence coordinates and the genomic regions without predicted genes.

*For a better understanding, the C3Q pipeline with the “Q” Cufflinks parameters, the subsampled BAM from RNA-Seq libraries for all the sequenced conditions (“ES3037M” for C. neoformans H99 and “ES3037” for C. deneoformans JEC21), the sequence filtering (sequences up to 150 nt, intronless sequences up to 300 nt, genome-predicted sequences without reads and alternative transcripts), and the Exonerate-based retrieval of deleted and non-predicted genes is named as C3Q6 protocol in the results section.*

### Automatization of the C3Q pipeline

The C3Q pipeline, an automatic gene predictor, was built with Python3 code *(C3Q_gene-predictor.py)* and is available in Github (https://github.com/UBTEC/C3Q)

The C3Q pipeline includes all established parameters for *Cryptococcus* genome annotation (C3Q6 protocol):

– The Cufflinks assembly of transcripts for each RNA-seq library.
– The merging of the generated GTF files by Cuffmerge.
– The GTF conversion to GFF format (needed for CodingQuarry).
– The training and genome prediction by CodingQuarry, using the merged GFF file and the reference genome.
– The sequence filtering: deletion of small transcripts up to 150 nt and intronless transcripts up to 300 nt; deletion of genome-predicted sequences without supporting reads and deletion of alternative transcripts from multi-transcript loci.
– The retrieval of deleted and non-predicted orthologous/paralogous sequences by Exonerate (modified version with GFF3 support from https://github.com/hotdogee/exonerate-gff3).

### Gene prediction in *C. deuterogattii* R265

Gene prediction in *C. deuterogattii* R265 was performed with the C3Q pipeline (C3Q6 protocol) using the *C3Q_gene-predictor.py* script. For this, the five RNA-seq triplicate libraries from *C. deuterogattii* R265 (Exponential phase at 30°C, Exponential phase at 37°C, Stationary phase at 30°C, Stationary phase at 37°C and Mating) were subsampled to 7.5 million reads each, and input into the script in addition to the *C. neoformans* H99 and *C. deneoformans* JEC21 protein sequences for the Exonerate step.

Concomitantly, manual correction of genes from chromosomes 9 and 14 was performed with the software Artemis (Carver et al., 2012), the R265 genome (NCBI assembly GCA_002954075.1), and the stranded paired-end RNA-seq data from *C. deuterogattii* R265 in the five growth conditions.

The predicted annotation was evaluated by comparing it to the manually corrected genes from chromosomes 9 and 14, as well as the *C. deuterogattii* R265 annotations from Broad (NCBI assembly GCA_000149475.3) and Ferrareze et al., 2017.

CDS gene coordinates from old annotations were also identified in the new sequenced genome with Exonerate aligner (*coding2genome*). The predicted novel genes were named with CNBG ID numbers above 10000. The statistics of the gene annotations ware generated by AGAT script *agat_sp_statistics.pl* (https://github.com/NBISweden/AGAT). The final annotation is available in file S1.

### Comparison of orthologue groups across *Cryptococcus* species

Ortho-groups and genes unique to *C. neoformans* H99, *C. deneoformans* JEC21 and *C. deuterogattii* R265 were evaluated with Orthofinder v2.3.3 configured to use the Blast aligner. Gene size comparisons were performed with orthologues and paralogues (if the true orthologue was not known) obtained from the OrthoFinder analysis (Emms and Kelly 2019), as well as gene sizes. For the ratio calculation, the size (nt) of the R265 gene was divided by the size (nt) of the H99 and JEC21 orthologous genes. The analysis of conserved domains in unique sequences and the functional annotation of *C. deuterogattii* R265 were performed with Blast2GO (Blastp, Interproscan and GO mapping).

### Gene orientation analysis

To determine the frequency of tandem genes with the same orientation, we searched for groups of two, three, four, or five genes assigned to the same DNA strand in the GFF file from *C. deuterogatti* R265 (with our new annotation), *C. neoformans* H99 (Genome assembly reference GCF_000149245.1), and *C. deneoformans* JEC21 (Genome assembly reference GCF_000091045.1) annotations. This was performed by analyzing the orientation of each gene pair in the GFF file from R265, H99 and JEC21, recording the frequency of genes converging (tail-to-tail), diverging (head-to-head), and in the same orientation (head-to-tail) in the whole genome and for each chromosome. This was executed using an in-house Python script (script *gene_organization_analysis.py*).

### Antisense transcription analysis

To evaluate the antisense transcription in the genomes analyzed, we first generated a reversed annotation, which consisted of a GFF file with genes assigned to the opposite strand of their actual annotation. With the annotation and the reverse annotation, we analyzed the percentage of antisense transcription for each protein coding gene using the software HTseq-count using the following attributes *(-f bam -r pos – s yes -t CDS -i ID -m intersection-nonempty --nonunique none (.bam) (.gff))* and the distinct RNA-seq libraries. Sense/antisense counts ratios for each gene for each condition were plotted. The script used for generation of a reverse GFF is available (*reverse_gff.py*).

### Intron retention evaluation

For a given intron, an IR index was calculated by determining the ratio of spliced:non-spliced reads at the upstream and downstream exon-intron junctions and choosing the lowest of these two numbers. These IR indices were calculated using RNA-seq obtained from cells growing in each of the four growth conditions. An intron was considered to be regulated by intron retention when the IR index was at least 0.01. We restricted our analysis to introns with more than 10 spliced reads.

### Statistical analysis

The proportion of genes with intron retention regulation was compared amongst the distinct conditions using one way ANOVA followed by multi-comparison analysis corrected by FDR. The X-squared analysis was conducted using R (version 4.0.2) and plots were prepared using the corrplot package (version 0.84).

### Availability and accession number

Raw and summarized sequencing data are available at SRA with the accession number: PRJNA660459. The C3Q pipeline is available in Github (https://github.com/UBTEC/C3Q); Supplemental files available at FigShare. The final annotation of *C. deuterogattii* genome was submitted to NCBI and is available on accessions CP025759.1 to CP025772.1.

## RESULTS

### Pipeline selection

We first compared the performance of two previously published annotation pipelines used in coding gene *de novo* annotation in intron-rich fungal genomes using RNA-seq data. The BRAKER1 pipeline, which combines GeneMark-ET (Lomsadze *et al.* 2014) and Augustus (Stanke *et al.* 2008) and is already optimized with the best prediction parameters (Hoff *et al.* 2016), was compared with a genome annotation pipeline composed of Cufflinks v2.1.1 (Trapnell *et al.* 2010) and CodingQuarry v2.0 (Testa *et al.* 2015). We used the *C. neoformans* H99 and *C. deneoformans* JEC21 genomes as controls to assess of the performance of both pipelines (Gonzalez-Hilarion *et al.* 2016; Wallace *et al.* 2020).

For this analysis, we used only RNA-seq data obtained in biological triplicate from cells grown to exponential growth phase at 30°C in complete medium (YPD) (EXPO30) (Wallace *et al.* 2020). Previously described BAM files obtained after alignment of trimmed reads to the *C. neoformans* H99 genome were input into the BRAKER1 and Cufflink-CodingQuarry pipelines (Wallace *et al.* 2020). For the Cufflink-CodingQuarry-based analyses, we used two alternative protocols. In the first case, we first merged the BAM files from each of the three replicates (CUFF-CQ protocol). In the second case, each replicate BAM file was used to generate a unique GTF prediction file, these files were then merged using Cuffmerge and used by CodingQuarry as a single transcript source (C3Q1 protocol) (Figure 1).

**Figure 1.**
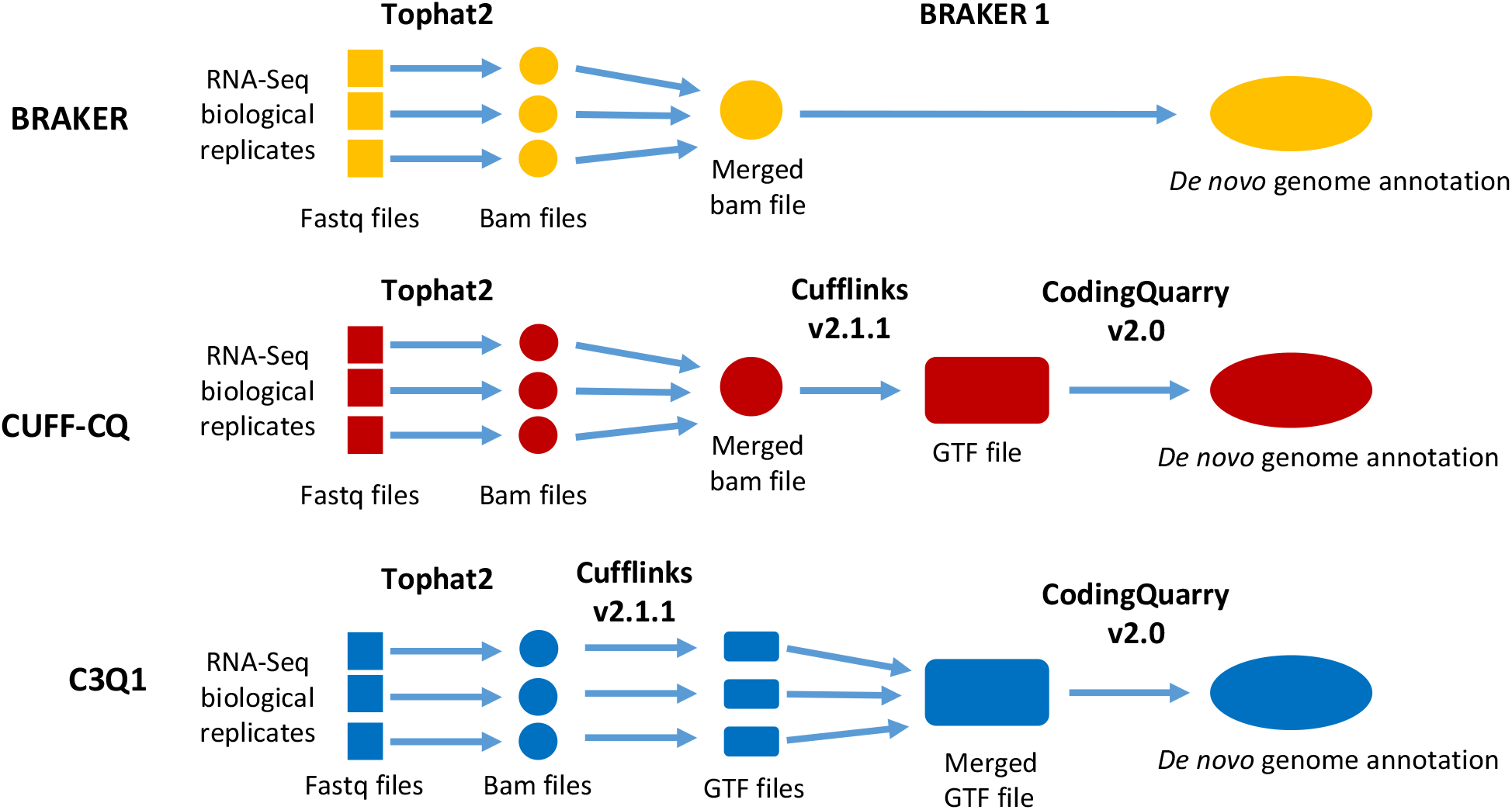
Schematic of the different pipelines tested in this study.

To compare the quality of these pipelines for identification of coding genes, we calculated their sensitivity (percentage of coding genes present in the reference annotation overlapping with one coding gene in the *de novo* annotation) and their specificity (percentage of predicted coding genes overlapping with one coding gene in the reference annotation). These comparisons revealed that BRAKER was much more sensitive than either Cufflinks-CodingQuarry protocol, missing only 91 coding genes in the *C. neoformans* genome (Figure 2A, Supplementary Table S1). However, the BRAKER pipeline was less specific (91.4%), predicting 622 coding genes absent in the reference annotation (Figure 2B). In contrast, both Cufflink-CodingQuarry protocols missed more coding genes (743 and 447 genes for CUFF-CQ and C3Q protocols, respectively), but had a higher (95%) specificity (Figures 2A,2B). We observed a similar pattern when we looked at CDS introns and CDS exons within the identified references genes. Again, the BRAKER pipeline was very sensitive, with only 0.4% missed introns (n=157) and 0.4% missed exons (n=164) in the prediction but had poor specificity, with 4471 novel introns and 3065 exons predicted but not present in the reference annotation (Figures 2A,B; Table S1). On the other hand, both Cufflink-CodingQuary-based protocols missed more introns (n=4944 and n=3238 for CUFF-CQ and C3Q1 protocols, respectively) and more exons (n=4281 and n=2777, respectively) but both were more specific, predicting less than 200 introns or exons not present in the reference annotation. Overall, both Cufflink-CodingQuary-based protocols returned more conservative results; they were more specific in the predicted gene structures and identified a smaller number of new insertions (novel exons/introns) and new genes. These more conservative predictions came at the cost of missing a larger number of features than the BRAKER protocol.

**Figure 2.**
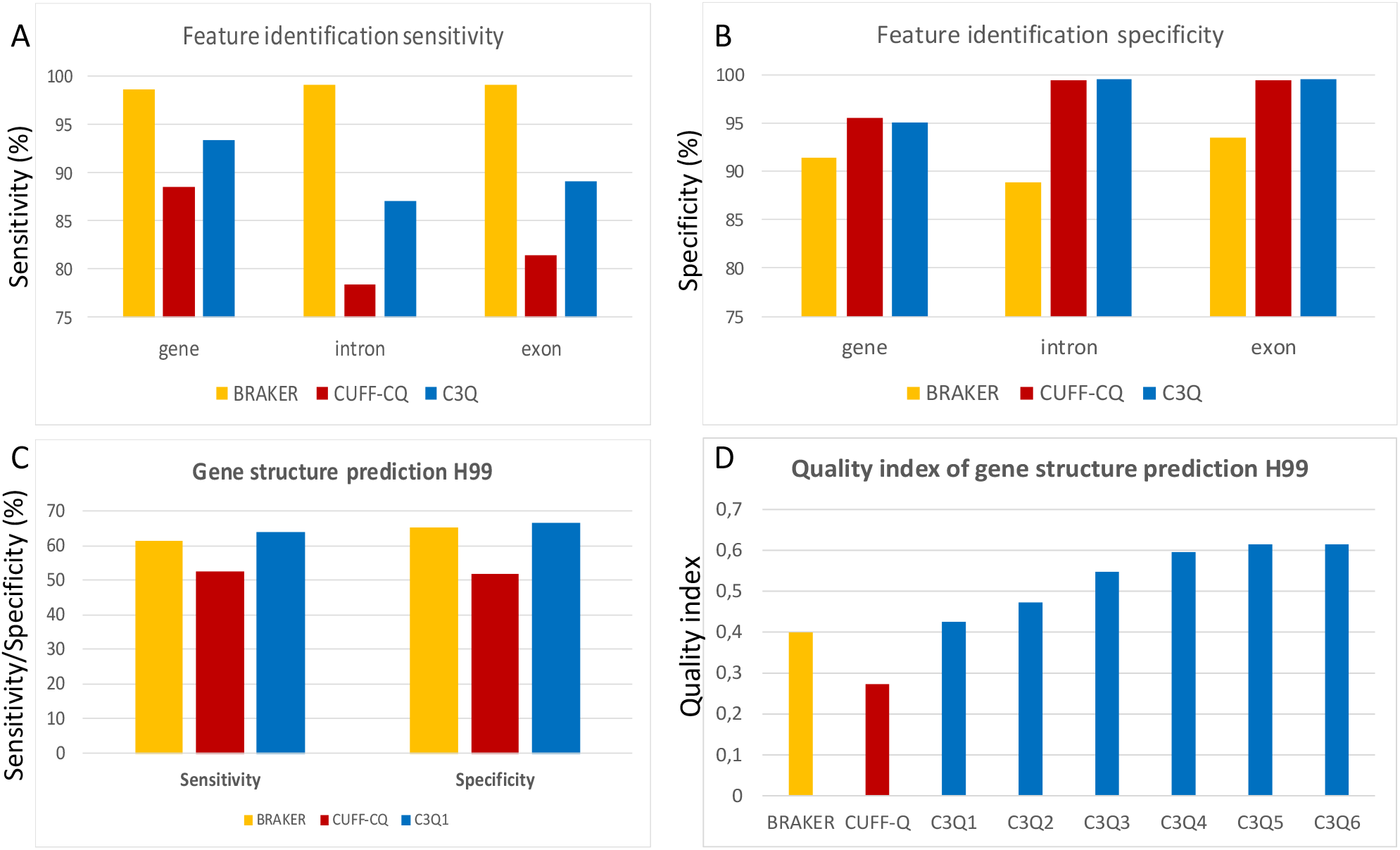
Sensitivity (A) and specificity (B) of the different tested pipelines for *C. neoformans* H99 genomic feature identification. For introns and exons, calculations were done using only genes that were both identified by the pipelines and present in the reference annotation. (C) Sensitivity and specificity of gene structure predictions using the three annotation pipelines. (D) Optimization of the C3Q pipeline. C3Q1 is the pipeline using default settings. C3Q2 through C3Q6 refer to the results obtained after each step of the pipeline optimization.

To assess all of these performance parameters and select the highest-performing protocol for further optimization, we considered the sensitivity and specificity of accurately predicting gene structure (perfect exon/intron organization) for each of the three pipelines. The C3Q1 protocol was the most sensitive, perfectly predicting the exon-intron layout of 66.5% (n=4516) of *C. neoformans* H99 genes, compared to 65.2% and 51.9% perfect predictions from the BRAKER and CUFF-CQ protocols, respectively (Figure 2C). This was also the most specific protocol with 63.9% of the predicted genes perfectly fitting the reference gene structures, compared to 61.3% and 52.6% of the predictions made by the BRAKER and CUFF-CQ protocols, respectively (Figure 2C). To better compare the quality of these pipelines, we considered a quality index that multiplied the sensitivity by the specificity of predicted gene structure predictions (Figure 2D). Our results demonstrated that the C3Q1 pipeline was the best, with a quality index of 0.42. We performed the same analysis with the *C. deneoformans* JEC21 genome annotation data and obtained similar results, confirming the C3Q1 protocol was the best protocol for further optimization (Figure S1).

### Optimization of the C3Q1 pipeline

#### Effect of Cufflinks settings

To improve both the number of perfectly predicted gene structures and the percentage of predicted loci in perfect agreement with the reference coding gene structures, we considered 17 combinations of Cufflinks settings. We varied parameters including 1) the minimum distance between transfags allowed to merge, 2) trimming of 3’ ends of reads, 3) filtering of alignments that lie within intronic intervals, 4) filtering of suspicious spliced reads, 5) minimum RNA-seq fragments allowed to assemble transfags, and 6) filtering of alignments that lie within intronic intervals in the same set of RNA-seq data. These Cufflinks parameter modifications reduced the number of missed genes and increased the number of reference genes identified from 6348 to 6462 (Table S1). Using the final settings, the pipeline C3Q2 quality index reached a score of 0.473, with 70.8% of reference gene intron-exon structures perfectly predicted and 66.8% of the predicted genes perfectly matching the reference exon-intron gene structures (Figure 2D).

#### Effect of the RNA-seq data set

We tested the C3Q2 optimized pipeline using additional RNA-seq data obtained under five different conditions in triplicate: stationary growth at 30°C (STAT30) and 37°C (STAT37), exponential growth at 30°C (EXPO30) and 37°C (EXPO37), and growth under mating conditions (Mating). Each RNA-seq data set generated a similar number of predicted transcripts, ranging between 7049 genes using the STAT37 set up to 7199 loci using the EXPO30 data set (Table S1). When compared to the reference set of genes, the number of predicted annotations were also very similar (Table S1). As expected, including more samples improved the annotation quality. The usage of the five conditions improved the C3Q3 annotation quality index to 0.547 despite the fact that more predicted genes not present in the reference genome were identified using this pipeline (n = 510) (Figures 2D, S2, Table S1). Similar results were obtained using the *C. deneoformans* JEC21 annotation and RNA-seq data (Figure S1, Table S1).

#### Evaluation of RNA-Seq data set size in gene prediction quality

Analysis of the results obtained using the C3Q2 pipeline fed with individual replicates of the EXPO30 RNA-seq data counterintuitively suggested the size of the initial BAM file might be negatively correlated with the quality of the final prediction (Tables S1). Identical analysis performed with *C. deneoformans* RNA-seq gave a similar result, suggesting the sequencing depth may substantially affect the quality of predictions and should be considered as a possible parameter of optimization for gene prediction pipelines. To improve the analysis of the effect of the size of the data set on the quality of gene prediction, the C3Q2 pipeline was tested with different representative fractions of reads from a single EXPO30 replicate. Thus, replicate samples with 1, 5, 7.5, 10, 15, 20, 30, and 40 million reads were used for *de novo* annotation of the *C. neoformans* H99 genome, and the quality of the gene predictions were compared. We performed this analysis using the same strategy for *C. deneoformans.* As shown in Figure 3, the quality of the gene structure prediction was highly dependent on the size of the RNA-seq initial data set in both species and strongly anti-correlated with the number of Cufflinks-assembled transcripts (Table S1). The highest-quality predictions were obtained with replicate samples with only 5-10 million reads. Using this adjusted read depth, the prediction showed improvement in nearly all parameters, including for missed genes, missed exons, and missed introns (Table S1).

**Figure 3.**
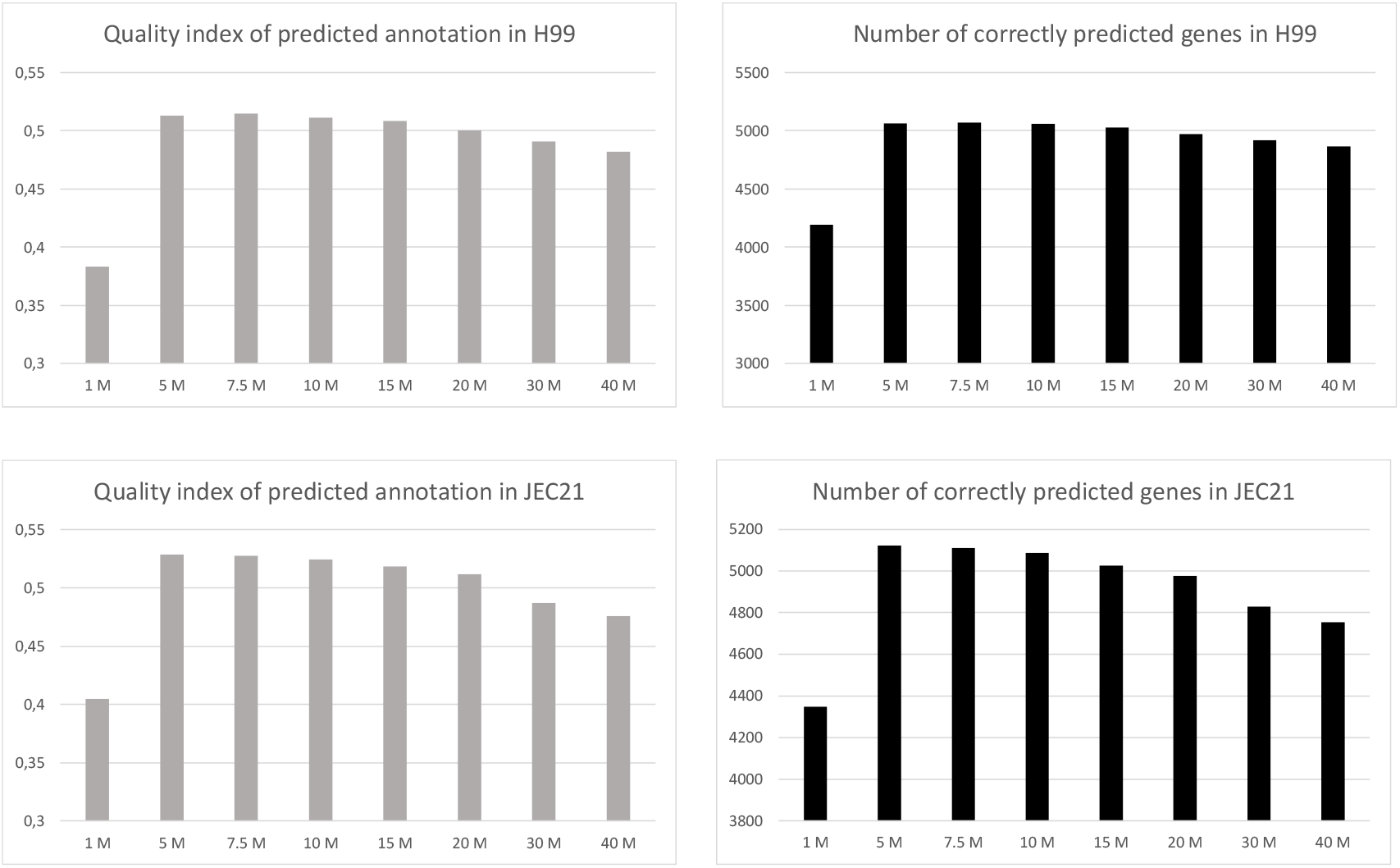
Effect of the size of the BAM file on the quality of the predicted annotation and on the number of correctly predicted genes in *C. neoformans* H99 and *C. deneoformans* JEC21.

We adjusted the number of reads to 7.5 million for each replicate in each condition and used these adjusted RNA-seq data sets for *de novo* annotation of the *C. neoformans* and *C. deneoformans* reference genomes. As expected, the gene predictions obtained with the C3Q4 pipeline were further improved with a quality index of 0.593 and 0.596, for *C. neoformans* and *C. deneoformans* annotations, respectively (Figure 2D, Figure S1, Table S1). In *C. neoformans*, 81.9% of the reference gene structure was perfectly predicted and only 1.9% (n=129) of genes were missed.

#### Gene filtering

Each optimization step improving the quality of the gene prediction was also associated with an increase of the number of predicted genes not present in the previously annotated reference genome (Table S1). Using the C3Q5 protocol, 717 (703 loci) and 774 (762 loci) additional genes were predicted in *C. neoformans* and *C. deneoformans*, respectively, compared to the reference annotation. The majority of these genes are likely to be misannotations. One hundred and six of the sequences had domain signatures of transposable elements, suggesting they correspond to fragments of transposons or retrotransposons unannotated in the reference H99 genome. To filter out some of the novel predicted genes, we looked at their structure and coverage. We compared the characteristics of these false-positive genes to the reference genes and found that most novel predicted genes were short (219 nt mean length, 112 nt median length), poorly expressed, and intronless. We tested different filters alone and in combination to eliminate as many false positive genes as possible without affecting the number of correctly predicted ones; the results are presented in Table S1. In both species, the best combination of filters eliminated all spliced coding regions smaller than 150 nt, all intronless genes smaller than 300 nt, and all genome-predicted genes not supported by any RNA-seq reads. Due to the presence of secondary transcripts at some loci, many of which were generated due to differences in the RNA-seq-predicted and genome-predicted transcripts for the same gene, a fourth filtering step was performed. In this step, to ensure that there was only one transcript per loci, the longest RNA-seq-predicted transcript or the longest genome-predicted transcript (for loci without RNA-seq evidence) was selected as a representative for the gene CDS coordinates. After this fourth filtering step, the number of predicted genes not present in the reference annotation was down to 409 and 427 genes in H99 and JEC21, respectively, and the quality index of the annotation increased to 0.614 and 0.608, respectively (Figures 2D, S1; Table S1).

#### Exonerate-based recovery of missed genes

Improvement of the pipelines was associated with an increase in the sensitivity of gene identification. In the initial C3Q1 protocol, 447 reference genes were missed, whereas only 162 H99 genes and 132 JEC21 genes were missed with the C3Q5 pipeline. Blast2GO analysis of the protein sequences encoded by the missed genes identified 16 proteins with conserved domains suggesting that it might be possible to identify some of them through comparative sequence analysis. Another 111 sequences were defined as hypothetical proteins. We first used the sequence alignment program Exonerate (Slater and Birney 2005) and the JEC21 reference proteome as a reference to try to recover missed coding genes in H99. As expected, this analysis identified a number of missed loci, but also added a number of unpredicted loci thus reducing the quality of the annotation. In the final C3Q6 pipeline, we chose to restrict this Exonerate analysis to genes that had been filtered out in the last step of the C3Q5 pipeline. We ultimately identified 14 and 9 novel genes in H99 and JEC21, respectively. Overall, the C3Q6 optimized pipeline was able to identify nearly 98% of genes in H99, contributing only 410 (~6%) novel genes. Importantly, the exon-intron structure of the predicted genes was predicted perfectly for >81% of the reference genes in both species.

### Intron number is predictive of the quality of the C3Q6 predicted annotation

During the course of the C3Q pipeline optimization, we obtained 88 versions of the H99 genome annotation. We carefully examined the different characteristics of these annotations, looking for a parameter predictive of their quality. First, we plotted the number of predicted loci against the quality of the annotation, but we did not observe any correlation. Similar results were obtained when we looked at missed or novel loci, suggesting that these parameters were also not indicative of the annotation quality. However, when the numbers of introns predicted were plotted against the quality of the annotation, we obtained a linear correlation (Figure 4). This correlation was lost during the filtration steps (red dots), which tend to reduce the number of introns. Similar results were obtained for JEC21, suggesting that the number of introns is a good parameter to consider when evaluating the quality of the annotation using the C3Q pipeline.

**Figure 4.**
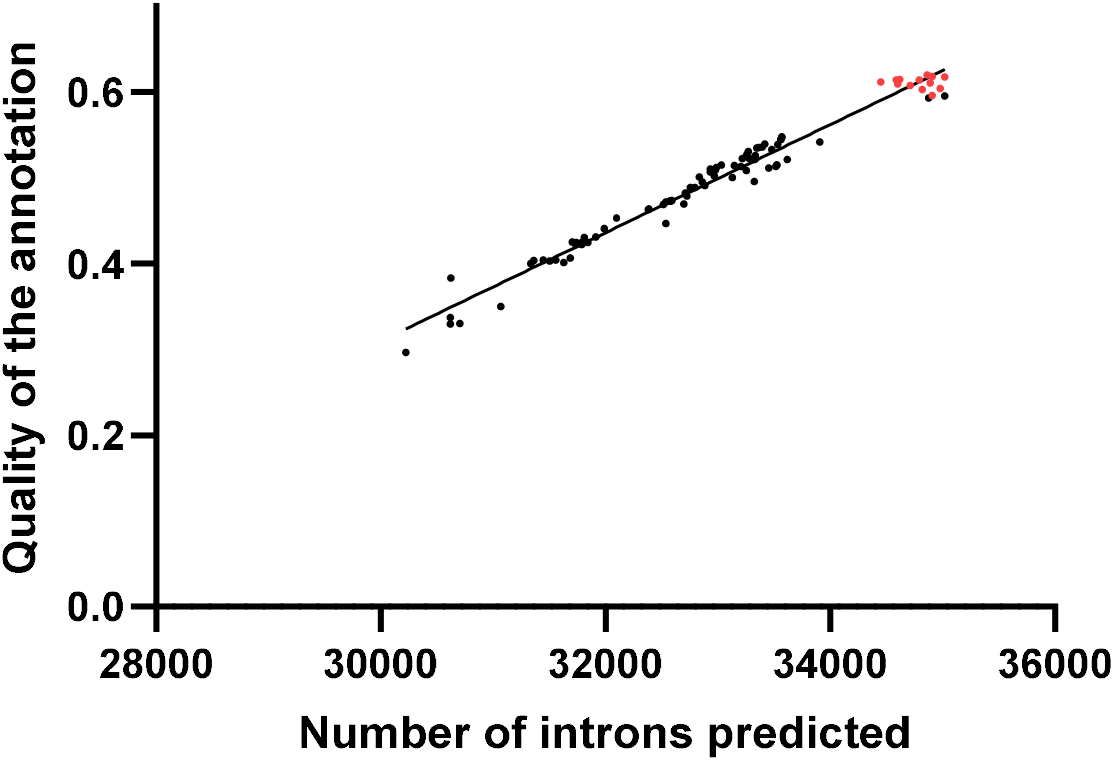
Relationship between the quality index of the H99 annotation and the number of introns and transcripts predicted. The red points correspond to the filtering steps of the optimization pipeline.

### Genome annotation of the *Cryptococcus deuterogattii genome*

We used the C3Q6 optimized pipeline to generate a new genome annotation for the *C. deuterogattii* reference strain R265. This reference strain was isolated in 2001 from the bronchoalveolar lavage fluid of an infected patient from the Vancouver Island outbreak (Kidd *et al.* 2004). Because of its outbreak origin and the loss of a functional RNAi pathway (D’Souza *et al.* 2011), *C. deuterogattii* has been the focus of a number of studies in recent years (Cheng *et al.* 2009; Ma *et al.* 2009; Ngamskulrungroj *et al.* 2012; Huston *et al.* 2013; Lam *et al.* 2019). The R265 genome has been previously annotated three times (D’Souza *et al.* 2011; Farrer *et al.* 2015; Ferrareze *et al.* 2017), but a recent release of telomere-to-telomere genome sequence data (Yadav *et al.* 2018) motivated us to generate an updated annotation. We generated RNA-seq data in biological triplicate from cells grown under five conditions (exponential growth phase at 30°C and 37°C, stationary growth phase at 30°C and 37°C, and under mating conditions) as previously described for *C. neoformans* H99 and JEC21 (Wallace *et al.* 2020). Reads were trimmed, aligned to the reference genomes (Table S2), and input into the optimized C3Q6 genome annotation pipeline.

To gain further insight into the quality of our updated R265 annotation, the structure predictions of genes for *C. deuterogattii* R265 chromosomes 9 and 14 were manually curated through visual examination of read alignments using Artemis (Carver *et al.* 2012) as previously described (Janbon *et al.* 2014; Gonzalez-Hilarion *et al.* 2016). We compared this manually curated annotation of chromosomes 9 and 14 with the prediction obtained from the C3Q6 genome annotation pipeline of these two chromosomes. This analysis revealed a quality index of this annotation of 0.51, with 68% of all predicted loci correctly predicted and 75% of the manually curated genes on these two chromosomes correctly predicted. In *C. neoformans* and *C. deneoformans*, the C3Q6 genome annotation pipeline missed very few genes (1.4% missed) and predicted a small number of false-positive genes (6.3%) (Table S3). As expected, the quality of the C3Q6 annotation was much better than previously published annotations (Farrer *et al.* 2015; Ferrareze *et al.* 2017) (Table S3).

### Manual curation of R265 annotation

To systematically analyze critical points of the automated *C. deuterogattii* R265 gene prediction, four sets of data were evaluated and selected for manual correction: 1) Exonerate-retrieved sequences (deleted and non-predicted), 2) predicted novel loci, 3) genes predicted in merged/split loci, and 4) small and potential pseudogenes. During the course of this manual curation of chromosomes 9 and 14, visual examination of the aligned reads revealed a number of loci at which the genome sequence did not entirely align with the RNA-seq reads, suggesting there were errors in the reference genome assembly. These errors were responsible for gene shortening or splitting and might partially explain the lower quality index score calculated for the R265 predicted annotation of chromosomes 14 and 9 compared to the quality scores obtained using similar data from JEC21 and H99. To systematically identify these types of annotation mistakes, we compared the size of the *C. deuterogattii* R265 predicted genes with their *C. neoformans* H99/*C. deneoformans* JEC21 orthologous counterparts. We identified 729 genes in R265 that were significantly smaller than their *C. neoformans* and *C. deneoformans* orthologues (size ratio < 0.8). Visual examination of these loci revealed that most of them were incorrectly predicted and needed manual curation. Manual curation was also performed for 67 genes that were significantly smaller than only one of their orthologues (*C. neoformans* or *C. deneoformans*). This manual curation also identified 125 genes which would have otherwise been challenging to predict due to genome sequence errors mistakenly affecting orthologue size ratios (Table S4).

Overall, our new version of R265 genome annotation contains 6,405 coding genes with 33,619 introns in CDS regions (34,512 introns including the UTRs). The manually corrected genes from chromosomes 9 and 14 were added and replaced the predicted genes from these regions, improving the quality of the final annotation. Of the 6,405 genes predicted with the C3Q6 pipeline, the CDS structure was modified for 873 coding through manual curation. Annotation of 3’UTR and/or the transcript leader sequence was performed for 1210 genes from the manually curated chromosomes (9 and 14) and the 873 manually curated genes with modified CDS structures. Furthermore, we annotated 55 lncRNAs and used tRNAscan (Lowe and Chan 2016) to annotate 161 tRNAs (Table S3). We also removed all genes predicted to reside within centromeric regions, and used previously published, manually curated annotations for these regions (Yadav *et al.* 2018).

### Putative pseudogenes and missing genes

We compared the gene content across the three annotated *Cryptococcus* genomes. We identified 5870 ortho-groups common to all three species (Figure 5). We found a similar number of R265-specific genes to the number of specific genes identified in H99 and JEC21, which is likely an indicator of the high quality of this annotation. Of interest, this analysis revealed 210 ortho-groups missing in *C. deuterogattii* R265, but present in both the *C. neoformans* H99 and the *C. deneoformans* JEC21 genomes (Table S5). This list of genes was curated first through Exonerate-based analysis and then through manual examination of the syntenic loci.

**Figure 5.**
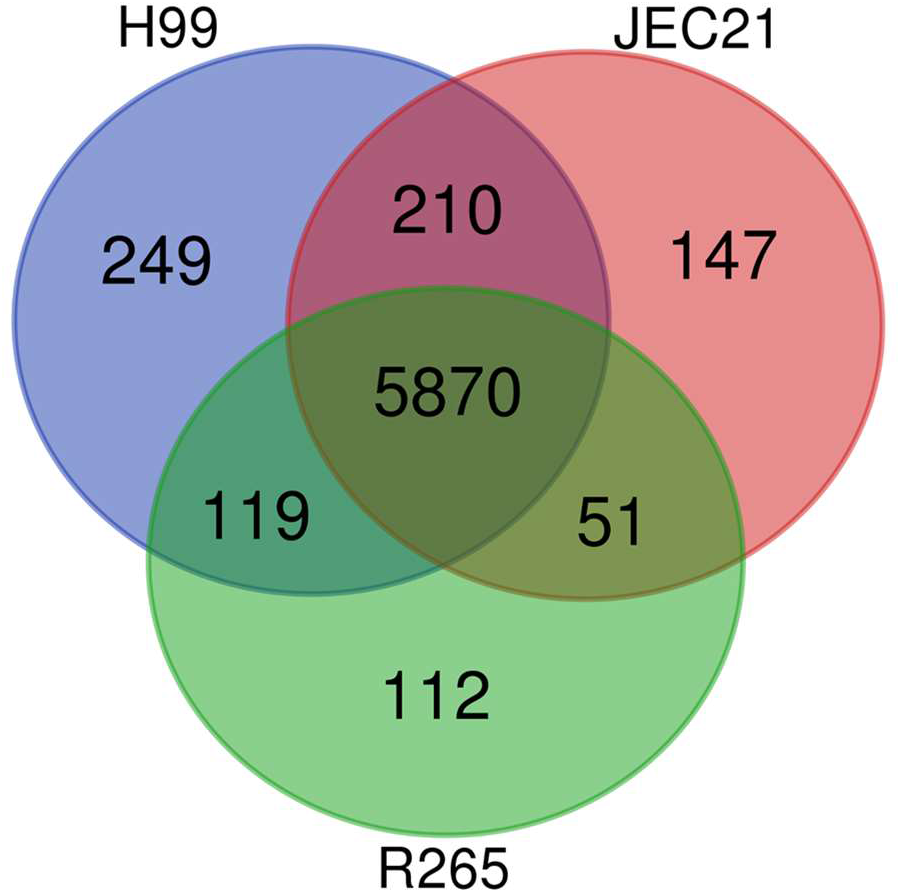
Comparative gene content of the annotated *C. neoformans* H99, *C. deneoformans* JEC21, and *C. deuterogattii* R265 genomes. Ortho-groups specific or common to the different species were identified and numbered.

*C. deuterogattii* R265 has previously been shown to lack a functional RNA interference pathway (D’Souza *et al.* 2011; Billmyre *et al.* 2013). Accordingly, the genes encoding one Dicer (*DCR1*) and an Argonaute protein (*AGO1*) have been lost, and the genes encoding an RNA dependent RNA polymerase gene (*RPD1*) and an RNAi essential zinc finger protein (*ZNF3*) are truncated and probably not functional in this strain (D’Souza *et al.* 2011; Feretzaki *et al.* 2016). The identification of truncated or absent genes in the R265 genome has been as a strategy to identify additional, novel components of the RNAi pathway in *C. neoformans* (Feretzaki *et al.* 2016). To identify genes specifically lost in *C. deuterogattii*, we considered the genes not predicted by our pipeline but present in the other *Cryptococcus* species annotations available in FungiDB (Basenko *et al.* 2018; Farrer ET AL. 2015; D’Souza ET AL. 2011). We considered here *C. tetragattii* strain IND107, *C. bacillisporus* strain CA187*3*, and *C. gattii* strains WM276, NT-10, and EJB2; no *C. decagatiii* annotation was available at the time of this study. We identified 17 ortho-groups that were absent in the R265 genome but present in all other species (Table 1). As expected, one ortho-group corresponds to an Argonaute protein (ortho-group OG0000415). We also confirmed that the gene *FCZ28,* which encoded a transcription factor essential for the sex-induced-silencing RNAi pathway in *C. neoformans*, was specifically absent in the R265 genome (Feretzaki *et al.* 2016). In contrast, the gene *GWO1*, previously identified as specifically lost in R265 and coding for an Ago1-interacting protein (although deletion mutants have normal siRNA profiles) was not present in this new list due to its absence in the IND107 genome (Dumesic *et al.* 2013). The orthologue size ratio analysis performed to pinpoint genome sequence mistakes eventually identified 119 R265 genes with a size ratio lower than 0.8 compared to both *C. neoformans* and *C. deneoformans* orthologs or shortened in one species and absent in the other (Tables 2; S7). Although some loci are likely pseudogenes, we decided not to annotate them as such because there is no strict structural definition of what constitutes a pseudogene (Tutar 2012) and we cannot evaluate the functionality of a gene based on its structure alone.

**Table 2.**
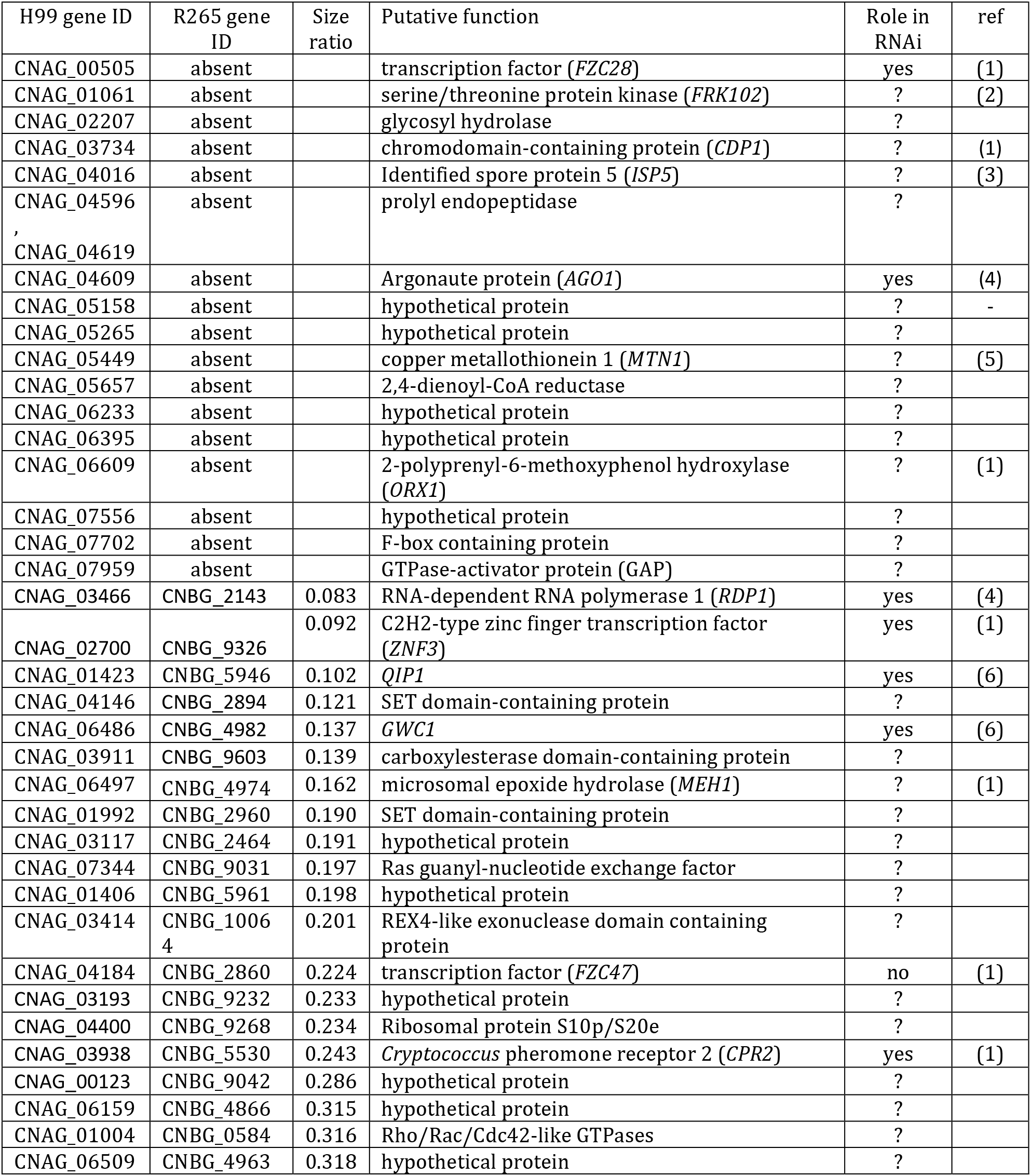
Genes with putative or known roles in RNAi identified as genes of H99 with orthologues in all *Cryptococcus* species but absent or severely truncated (and thus putative pseudogenes) in *C. deuterogattii* R265 as compared to JEC21 and H99 (proteins with a ratio <0.33 are presented. The full table of shortened genes is presented as Table S6). (1) Feretzaki et al 2016; (2) Lee et al. 2016; (3) Huang et al. 2015; (4) Wang et al. 2010; (5) Ding et al. 2011; (6) Dumesic et al. 2013.

As expected, the RNAi genes *RPD1 (Wang et al. 2010)*, *ZNF3 (Feretzaki et al. 2016)*, *CPR2 (Feretzaki et al. 2016)*, *QIP1 (Dumesic et al. 2013)*, *GWC1 (Dumesic et al. 2013)*, *RDE4,* and *RDE5* (Burke *et al.* 2019) were present in this list, confirming that a large number RNAi genes are not functional or are absent in R265. Conversely, *RDE1*, *RDE2,* and *RDE3* (Burke *et al.* 2019), which were recently implicated in RNAi in *C. neoformans*, all possess an orthologue of similar size in R265 (CNBG_3369, CNBG_4718, and CNBG_1922, respectively). Of note, in this version of the R265 annotation, the *DMT5* (CNBG_3156) gene encoding a putative DNA methyltransferase is not truncated as previously published (Yadav *et al.* 2018; Catania *et al.* 2020) and appears to be expressed and functional.

### Gene organization, antisense transcription, and alternative splicing in R265

The absence of RNAi in R265 was recently shown to be associated with a modification of the chromosome structure: shorter centromeres and the loss of any full-length transposable elements (Yadav *et al.* 2018). Here, we examined the possible consequences of RNAi loss on the transcriptome aside from the expected absence of siRNA. We first hypothesized that the absence of RNAi in *C. deuterogattii* could result in increased antisense transcription, as it might be the source of double-stranded RNA; increased antisense transcription in RNAi-deficient species has also been observed in *Saccharomyce*s species (Alcid and Tsukiyama 2016). We thus evaluated the sense/antisense transcript ratio at coding gene loci. Because the 3’UTR and TL sequences were only partially annotated in the R265 genome, we restricted our analysis to the CDS regions. We compared the ratio of read numbers of sense and antisense transcripts corresponding to all coding regions in *C. neoformans*, *C. deneoformans,* and *C. deuterogattii* under four growth conditions. When cells were grown to exponential growth phase at 30°C (E30), most of the expressed genes (92.2 %) had antisense transcription in *C. neoformans*, but antisense transcripts were expressed at a very low level (1.2% of transcription antisense *vs* sense). Antisense transcription prevalence and expression levels were similar in the two other species (92.6% and 95.2% of genes with antisense transcription, 3.2% and 2.5% of antisense *vs* sense transcription in *C. deneoformans* and *C. deuterogattii,* respectively). These ratios changed in different growth conditions, particularly increased temperature at both exponential and stationary growth phase. However, this analysis did not provide evidence of a link between the level of antisense transcription and the absence of RNAi in R265 because the RNAi-proficient JEC21 strain had the highest antisense/sense transcription ratio across all conditions tested (Figure 6).

**Figure 6.**
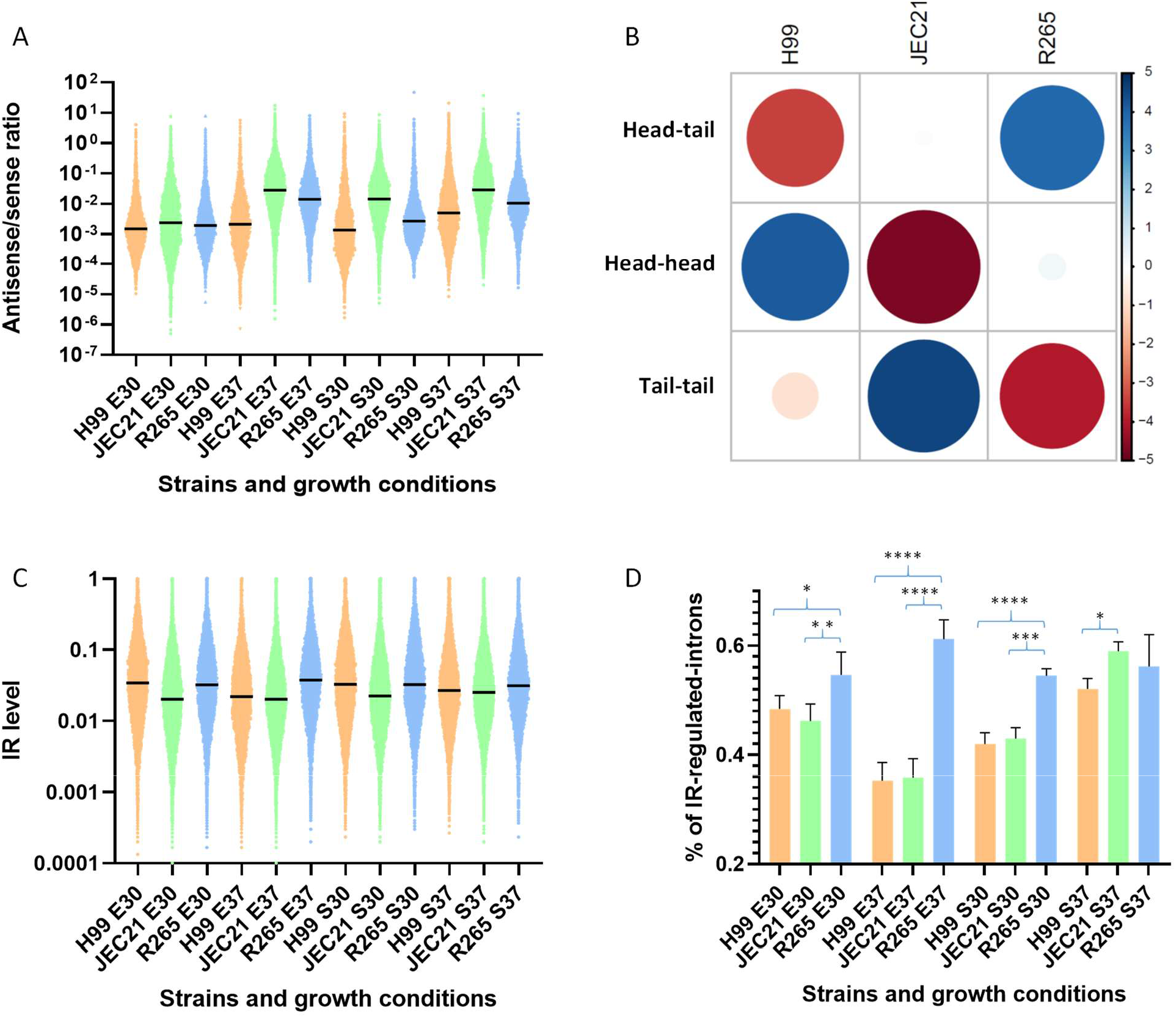
(A) Antisense/sense transcription ratios in *C. neoformans* (H99), *C. deneoformans* (JEC21) and *C. deuterogattii* (R265). RNA-seq data obtained from cells grown to exponential phase at 30°C (E30) and 37°C (E37) or stationary phase at 30°C (S30) and 37°C (S37) were used. (B) Statistical analysis (Pearson’s Chi-squared test) revealed a species-specific bias in gene orientation. Circle size is proportional to the standardized residuals, with absolute values higher than 2 representing statistical significance (Sharpe 2015). Positive values (blue circles) in cells specify a positive association between the corresponding row and column variables. Negative residuals are represented by red circles. This implies a negative association between the corresponding row and column variables. (C) Intron retention level in each species according to growth condition. Black bars represent median values. (D) Percentage of CDS introns regulated by IR in each species according to growth condition. The results of the statistical analysis (ANOVA one-way multi-comparison analysis corrected by FDR). * (q value < 0.05), ** (q value < 0.01), *** q value <0.001), **** (q value < 0.0001).

We then analyzed gene orientation in the three species, evaluating the number of genes coupled in a tail-to-tail orientation as this orientation should favor antisense transcription over head-to-tail or head-to-head orientations. Indeed, as shown in Figure 6B, there was a clear selection against tail-to-tail gene orientation in *C. deuterogattii*, thus limiting antisense transcription (c-squared = 103.79, df = 4, p-value < 2.2e-16). In contrast, this orientation is favored in JEC21, which might explain the higher level of antisense transcription.

The SCANR model predicts that siRNAs are produced in response to poorly spliced introns that stall the spliceosome complex, which should result in lower levels of expression for the corresponding gene (Dumesic *et al.* 2013). To explore whether loss of RNAi could have affected intron retention (IR) in *C. deuterogatiii*, we compared the number of CDS introns regulated by intron retention in this species and two RNAi-proficient ones. Interestingly, in three conditions the percentage of introns regulated by IR was higher in R265 than in JEC21 or H99 (Figure 6C). For instance, when cells were grown to exponential phase at 37°C, 44.5% of R265 introns are regulated by IR as compared to 21.7% and 20.8% in *C. neoformans* and *C. deneoformans*, respectively. In contrast, the IR indices were similar across the three species when cells were grown at 30°C. However, at 37°C in either exponential and stationary growth phase, the median value of the IR index in R265 was higher than those in both *C. neoformans* and *C. deneoformans*. Overall, these data suggest that IR is better tolerated in R265 than in H99 or JEC21; this result aligns with the SCANR model of siRNA production and gene regulation in *Cryptococcus*.

### Subtelomeric gene organization and cluster exchange in *Cryptococcus*

Our analysis identified 210 orthologue groups present in both *C. neoformans* and *C. deneoformans* but absent in *C. deuterogattii*. Interestingly some of these lost genes are clustered in these genomes. We identified 21 clusters of lost genes with consecutive elements in both *C. neoformans* and *C. deneoformans* reference genomes. One of these lost clusters has been previously described and has been reported to contain homologues of several *GAL* genes (*GAL1*, *UGE2*, and *GAL7*) and a gene encoding a sugar transporter of the major facilitator superfamily (MFS) (CNAG_07897) (Slot and Rokas 2010). We also identified a fifth gene in this cluster (CNAG_06055) which encodes a putative alpha-1,4-galactosidase (Figure S2A). *C. neoformans* and *C. deneoformans* also possess unclusterered paralogues of the genes *UGE2* (*UGE1*, CNAG00697), *GAL1* (*GAL101*, CNAG_03946), and *GAL7* (*GAL701*, CNAG_03875). Previous studies have shown *UGE2* is required for growth on galactose, whereas it paralogous gene *UGE1* is necessary for growth at 37°C and glucuronoxylomannogalactan (GXMGal) biosynthesis, which makes up an important fraction of the *Cryptococcus* polysaccharide capsule (Moyrand *et al.* 2008). Interestingly, we previously reported that a *uge2Δ* mutant strain was able to grow on galactose at 37°C, suggesting that *UGE1* is able to compensate in the absence of *UGE2* at 37°C. The GAL cluster with five genes has also been lost in all other *Cryptococcus* species that were assessed. Thus, the *C. gattii* clade species possess the only non-clustered paralogues of the GAL pathway; yet, they are all able to grow on galactose as a sole carbon source, suggesting these genes are involved in both GXMGal synthesis and galactose assimilation in this species (Figure S2B).

Gene ontology (GO) term enrichment analysis (Priebe *et al.* 2014) of 52 genes within 18 non-subtelomeric clusters that were absent in R265 revealed a statistically significant enrichment of genes coding for proteins implicated in transport and transcription regulation (Figure 7A). Functional annotation of these genes confirmed this result (Table S7). We identified 13 clusters containing at least one gene coding for a putative transporter, including six MFS-type transporters, and eight clusters containing at least one gene coding for an annotated or putative transcription factor (TF), including six fungal Zn(2)-Cys(6) binuclear cluster domain-containing TFs. Overall, seven clusters contain both a transporter and a TF (Figure 7B, Figure S3). Strikingly, this association between transporters and TFs resembles the organization of primary metabolic gene clusters (MGCs) (Rokas *et al.* 2018). Because three MGCs were located within subtelomeric loci, we compared the gene content within subtelomeric regions to the gene content of the lost clusters. We considered the 20 most distal genes of each chromosome arm in H99. GO term enrichment analysis of these 560 H99 subtelomeric genes revealed very similar profiles to the profiles obtained for the cluster genes. Again, genes coding for proteins implicated in transport in subtelomeric regions were significantly enriched (Figure 7C).

**Figure 7.**
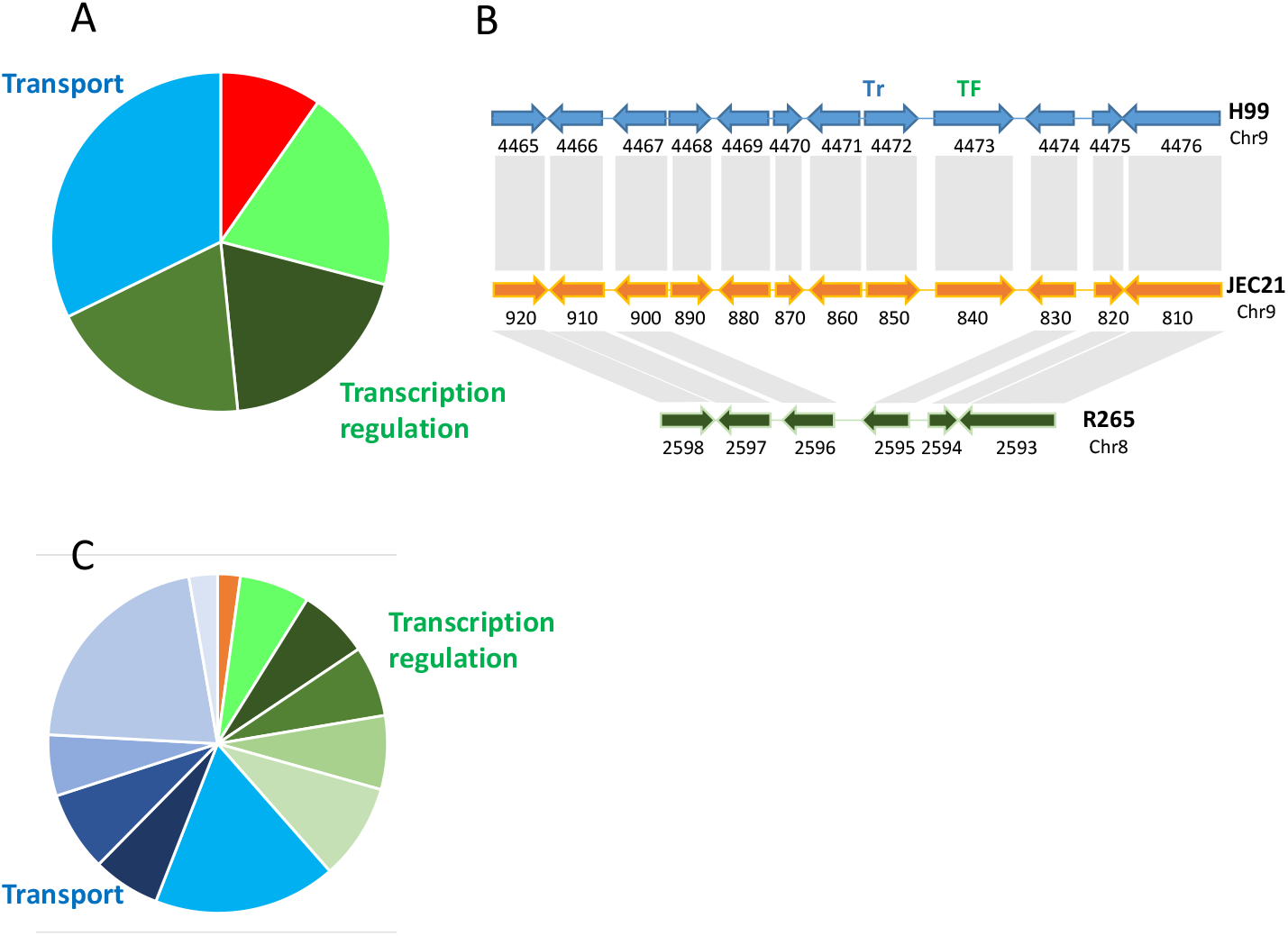
(A) GO term enrichment analysis of genes in clusters absent in R265. Green colors indicate GO terms associated with transcription regulation (GO:0006012, GO:0000981, GO:0006366, GO:0006357). Blue colors indicate GO terms associated with transport (GO:0055085). Orange colors indicate GO terms associated with galactose metabolism (GO:0006012). (B) Example of the organization of an MGC-like cluster absent in R265. CNAG_04468 (CNI00890) encodes a putative tartrate dehydrogenase, CNAG_04469 (CNI99880) encodes a putative 4-aminobutyrate transaminase, CNAG_04470 (CNI00870) encodes a putative halo-acid dehalogenase, CNAG_04471 (CNI00860) encodes an FAD-dependent oxidoreductase superfamily protein, CNAG_04472 (CNI00850) encodes an MFS protein, and CNAG_04473 (CNI00840) encodes a TF with a fungal Zn(2)-Cys(6) binuclear cluster domain. (C) GO term enrichment analysis of subtelomeric genes in H99. Green colors indicate GO terms associated with transcription regulation (GO:0051213, GO:0000981, GO:0006366, GO:0006357, GO:0006351, GO:0006355). Blue colors indicate GO terms associated with transport (GO:0055085, GO:0022891, GO:0022857, GO:0005215, GO:0016021, GO:0008643, GO:0006810). The orange color indicates a GO term associated with dioxygenase activity (GO:0051213).

Functional annotation of these subtelomeric genes confirmed this enrichment of transporters and TFs (Table S8). We found an unexpected number of genes encoding annotated or putative TFs (n = 33) and transporters (n = 68) within these regions of the H99 genome. Most of these TFs and transporters belong to the fungal Zn(2)-Cys(6) binuclear cluster domain-type (n= 24) and MFS-type (n= 49) families, respectively. Comparison of the organization of *C. neoformans* H99 subtelomeric loci to those in *C. deneoformans* JEC21 revealed a very similar organization, and only four mosaic subtelomeric regions were identified with genes from at least two different regions in H99; few genes were present in H99 but absent in JEC21 (Figure 8). However, we did identify two duplicated regions in the JEC21 subtelomeric regions. The first duplicated locus consists of six genes with orthologues in subtelomeric region of the left arm of Chr 5 in H99. The second duplicated region has been previously described (Fraser *et al.* 2005). It is located in the left arms of Chrs 8 and 12 and resulted from a telomere-telomere fusion that occurred during the construction of the JEC20/JEC21 congenic mating pair. Interestingly, a TF with a fungal Zn(2)-Cys(6) binuclear cluster domain (*FZC2*, CNAG_05255) and a putative amino acid transporter (CNAG_05254) are present within this repeated region. Conversely, genes in the subtelomeric regions of the right arms of H99 Chrs 4 and 14 are orthologues of genes located within a central part of JEC21 Chr 8 (Figure S3), suggesting an additional telomere-telomere fusion event. In contrast, the subtelomeric regions in R265 have undergone more rearrangements compared to JEC21 – in R265 there are fifteen mosaic subtelomeric regions that contain genes from at least two different regions in H99. We also identified nine genes within six R265 subtelomeric regions whose orthologues are located far from the telomeres in H99. Interestingly, functional annotation of the R265-specific subtelomeric gene clusters (n = 12) (Figure 8; Table S9), revealed an enrichment of genes encoding TFs (n = 2) and transporters (n = 6).

**Figure 8.**
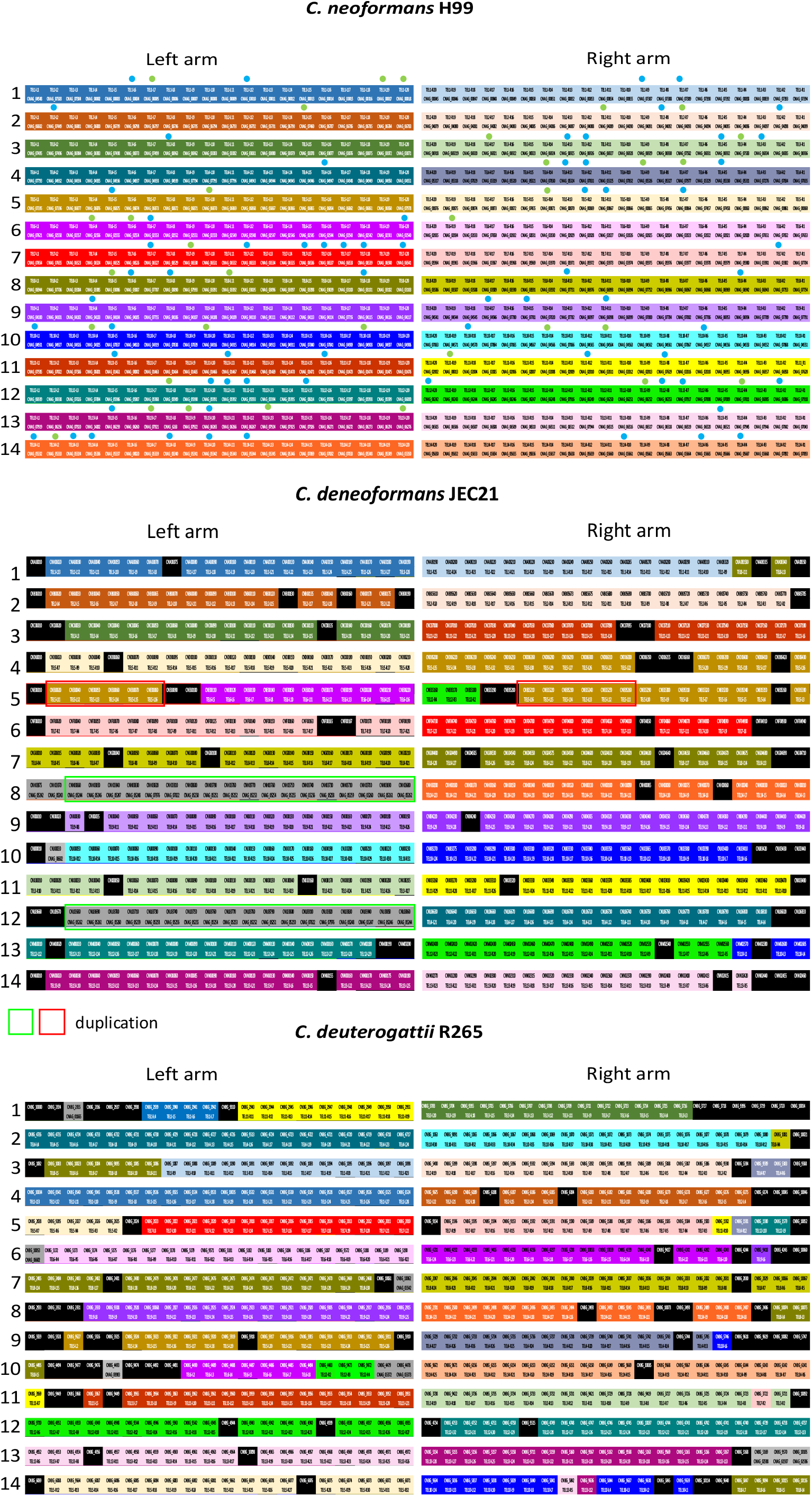
Subtelomeric gene organization in *Cryptococcus*. The 20 most distal genes at each subtelomeric locus were considered. The color code identifies each subtelomeric regions in H99 and orthologous genes in the other species. The positions of these orthologues in the H99 subtelomeric regions are given (TEL-RX or TEL-LX correspond to genes positioned within the right or left arm of chromosome X). When the orthologous gene is not located within a subtelomeric region, its locus named is given. Black boxes correspond to genes present in *C. deneoformans* or *C. deuterogattii* but absent in *C. neoformans*. Red and green boxes indicate duplicated sets of genes. Blue dots indicate transporters. Green dots indicate transcription factors.

Subtelomeric regions have been shown to be silenced by H3K27me3 histone modifications in *C. neoformans*, and a large number of genes that are upregulated upon deletion of the H3K27 methyltransferase (encoded by *EZH2*) are located within subtelomeric regions (Dumesic *et al.* 2015). Accordingly, we observed that expression of the 580 most proximal genes was generally lower than the expression of the most telomere-distal genes (Figure 9). Interestingly, we found that the H99 genes present within the MGCs that were lost in R265 were also poorly expressed. However, none of these genes were upregulated upon *EZH2* deletion (Dumesic *et al.* 2015), suggesting that they are not directly regulated by H3K7me3. In summary, these data suggest that dynamic exchanges of MGCs between subtelomeric regions occurred during *Cryptococcus* speciation. These results also suggest that MGC exchanges between subtelomeric loci and more central parts of chromosomes might be associated with new assimilation capacities.

**Figure 9.**
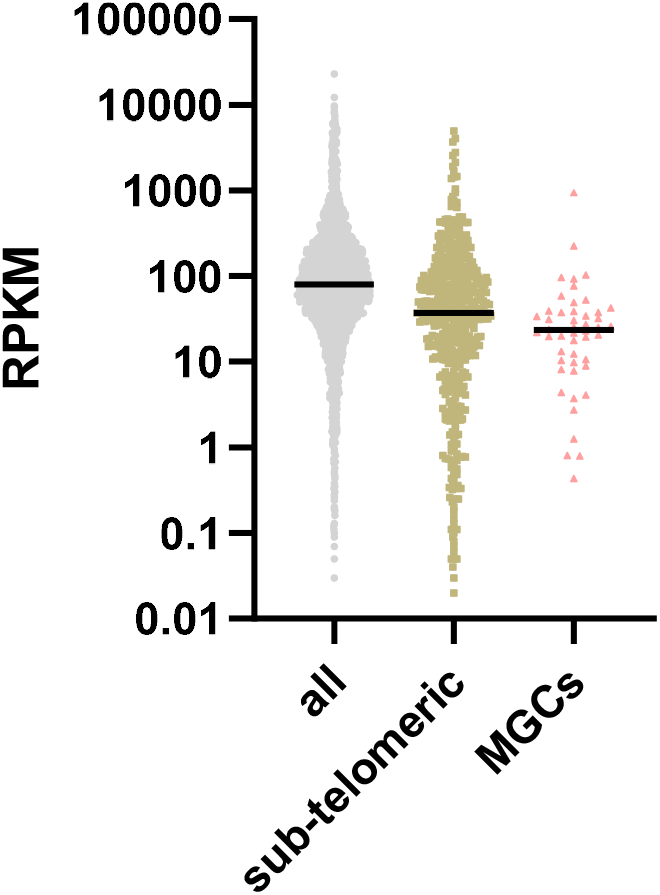
Expression of genes according to their position on the chromosome. Subtelomeric genes are the 20 most distal genes of each chromosome arm. The H99 genes present within the non-subtelomeric cluster of genes lost in R265 are indicated as MGCs.

## Discussion

Although a number of bioinformatic pipelines have been published in recent years, accurate annotation of fungal genomes is still difficult due to their complexity and compactness (Haas *et al.* 2011). In this study, we have carefully optimized a previously published Cufflinks-CodingQuarry-based annotation pipeline and tested it on two complex fungal genomes. This pipeline largely outcompeted the BRAKER1 pipeline when applied to two *Cryptococcus* reference genomes (*C. neoformans* H99 and *C. deneoformans* JEC21) and would likely outcompete many other pipelines use to annotate fungal genomes *de novo* (Min *et al.* 2017).

Our optimization process revealed three notable points. First and counterintuitively, increasing the quantity of data did not always result in a better annotation. This is likely because Cufflinks tends to make huge clusters when large data sets are input; these clusters might be eliminated during the transcript identification step. Accordingly, we found that the number of predicted transcripts decreased when too much data was used. Second, we found a nearly linear relationship between the number of introns predicted and the quality of the annotation. However, this correlation did not hold when two of the pipelines were compared; the BRAKER pipeline predicted more introns than the C3Q pipeline, along with predicting many more genes. Nevertheless, the correlation between intron number and annotation quality provided a facile way to evaluate the reliability of a *de novo* annotation, which might be affected, for instance, by the quality of the RNA-seq data. Third, we found the final step of comparative genomics did not always improve the quality of the annotation. In our assay, the Exonerate-based analysis step using the whole proteome of a reference species primarily introduced errors into the annotation. This was probably due to the fact that even when manually curating genome annotations, a number of dubious genes remain, which are then transferred to the new genome annotation. In fact, a systematic usage of a comparative annotation step following a *de novo* RNA-seq annotation would likely result in a dramatic expansion of dubiously annotated genes in fungal genomes. Accordingly, it is noticeable that the number of predicted coding genes in R265 (n=6405) is lower than the ones predicted in H99 (n=6795) and JEC21 (n=6639) although we ignore whether these differences have some biological relevance or if they are due to the different strategies used to annotate these genomes.

During the annotation of the R265 genome, we manually curated a subset of genes that were lost in R265 compared to all of the other *Cryptococcus* species as well as a set of putative pseudogenes. The identification of genes specifically lost or pseudogenized in R265 has previously been used as a strategy to identify novel RNAi components in *C. neoformans* (Feretzaki *et al.* 2016). Accordingly, most known RNAi genes are present in these sets of lost and pseudogenized genes (Billmyre *et al.* 2013). However, some genes, like *RDE1* (Burke *et al.* 2019), which is necessary for siRNA production, are present and functional in R265, suggesting that it may have roles independent from RNAi silencing. On the other hand, *GWO1*, which is also considered to be an RNAi pathway component, is also absent in the *C. tetragattii* strain IND107 and is therefore absent in our list as well. One possible explanation is that Gwo1 alone or in complex with Ago1 could play another role independent of RNAi. Another possibility is that a Gwo1-dependent RNAi pathway has also been lost in *C. tetragattii*. Nevertheless, this analysis confirms that looking for specific gene loss in a fungal species deficient for a certain pathway remains a promising strategy for the identification of genes implicated in this pathway in other proficient species. In the present case, it would be interesting to see how many of the R265 truncated genes are functional in other *C. gattii* species, although it would demand a complex manual curation, which is beyond the scope of this paper.

Our study revealed that loss of RNAi in R265 is associated with few general transcriptome modifications compared to the transcriptomes of JEC21 and H99, aside from the predictable absence of siRNA. This might be because we did not annotate most non-coding features like lncRNAs, transcript leaders, and 3’UTRs. Yet, quantification of the sense/antisense transcription ratio at CDS did not reveal any differences between R265 and the other *Cryptococcus* species analyzed, suggesting that this ratio does not depend on the RNAi status of the species in this genus. This is in agreement with the fact that siRNAs in *C. neoformans* primarily target transposons and retrotransposons (Janbon *et al.* 2010; Wang *et al.* 2010; Dumesic *et al.* 2013), whereas antisense transcription is associated with nearly all of the genes as we have shown. This result also suggests antisense transcription in *Cryptococcus* only rarely results in the production of double-stranded RNA. Dumesic and colleagues showed that delayed splicing is a source of siRNA production in *C. neoformans* (Dumesic *et al.* 2013). We thus anticipated that the absence of RNAi would increase the level of intron retention. In agreement with previous reports in *C. deneoformans*, we found that IR level was regulated by growth conditions in both *C. neoformans* and *C. deuterogattii* (Gonzalez-Hilarion *et al.* 2016). However, the number of introns regulated by IR was markedly larger in R265 suggesting that IR is better tolerated in this RNAi-deficient species. We also expected that some compensatory mechanisms might be acting to control the level of IR because IR rates were largely similar across the three species analyzed even though it was higher in R265, at least at 37°C. It is important to note the remarkable effect of temperature on both IR and antisense transcription, which might be related to a recent report that transposons are specifically mobilized at this temperature in *C. deneoformans* (Gusa *et al.* 2020).

While most loci that have been lost in R265 compared to other *Cryptococcus* species contain only a single gene, we also identified gene clusters that were missing in R265. Analysis of the gene content within these clusters revealed a strong enrichment of genes coding for proteins implicated in transport and transcriptional regulation. This finding was reminiscent of patterns identified in metabolic gene clusters (MGCs) involved in primary metabolism, which typically contain transcription factors and transporters (Rokas *et al.* 2018). MGCs can be lost or gained in fungi and several examples of instances of horizontal transfer of whole MGCs from one species to another have been published (Slot and Rokas 2010; Rokas *et al.* 2018; Wang *et al.* 2019). In filamentous fungi, the majority of MGCs are located within subtelomeric regions, which are largely subjected to inter-chromosomal reshuffling (Gladieux *et al.* 2014). Two examples of lineage-specific gene clusters harboring both transcription factors and transporters have been previously reported in *C. neoformans* (Rhodes *et al.* 2017), suggesting dynamic gene cluster gain and loss events even with a single species in *Cryptococcus*. Interestingly, these *C. neoformans* lineage-specific clusters also contain transcription factors and transporters (Rhodes *et al.* 2017). A more recent report suggests that genes within one of *these C. neoformans* clusters are co-regulated, as is expected from a typical MGC (Yu *et al.* 2020). In *Cryptococcus*, we found that the subtelomeric regions were also enriched for characteristic MGC genes as well, and comparisons of subtelomeric gene organization across the three *Cryptococcus* species suggested active reshuffling. This was in agreement with previous data showing that subtelomeric genes are under strong evolutionary pressure in *Cryptococcus* (Desjardins *et al.* 2017). We found a large number of genes encoding transporters and TFs of unknown function in *Cryptococcus* subtelomeric regions. Surprisingly, most of the TF genes identified in these MGCs within subtelomeric regions as well as in MGCs far from telomeres were not annotated as TFs and were not included when a systematic TF deletion collection was constructed and studied (Jung *et al.* 2015). It therefore seems that the TF repertoire in *Cryptococcus* may be larger than currently appreciated. Similarly, besides myo-inositol transporters, which have been previously reported to be localized within subtelomeric regions (Xue *et al.* 2010), the substrates of most transporters located in these regions remain unknown.

Genes within subtelomeric regions are silenced by H3K27me3 epigenetic marks and, accordingly, are expressed at lower levels than genes located in more central regions of the chromosomes. Similarly, genes within the subtelomeric clusters lost in R265 were poorly expressed. Yet, their expression levels did not significantly change following deletion of the gene encoding the H3K27me3 methyltransferase *EZH2,* suggesting they are either not regulated by H3K27me3 or that additional changes are needed to activate expression of these genes like those previously described in *Fusarium graminearum* (Connolly *et al.* 2013). If this is the case, the regulation of *GAL* genes by galactose might represent a good example of how genes within the MGCs could be regulated in *Cryptococcus* (Wickes and Edman 1995; Moyrand *et al.* 2008; Ruff *et al.* 2009). Besides the *GAL* cluster, the function and regulation of most of MGC genes in *Cryptococcus* are unknown. Nevertheless, our results suggest active exchange between subtelomeric regions and more central parts of chromosomes in *Cryptococcus*, potentially reshaping primary metabolism for adaptation to different environmental niches. They also emphasize how both complete genome and precise annotations are needed to study these dynamics in fungi.

## Acknowledgements

PAGF was supported a CAPES exchange grant (Advanced Network of Computational Biology RABICÓ – Grant no. 23038.010041/2013-13). This work was supported by a CAPES COFECUB grant n°39712ZK to GJ, by a CNPq grant (309897/2017-3) to CCS, and by a NIH/NIAID R37 MERIT Award AI39115-23 and a NIH/NIAID R01 Award AI50113-16 to J.H. J.H. is co-director and fellow of CIFAR program Fungal Kingdom: Threats & Opportunities. S.J.P was supported by the NIH/NIAID F31 Fellowship 1F31AI143136-02. Members of the Heitman laboratory are acknowledged for valuable discussion.

