## Supplementary material for "Application of an Optimized Annotation Pipeline to the *Cryptococcus Deuterogattii* Genome Reveals Dynamic Primary Metabolic Gene Clusters and Genomic Impact of RNAi Loss": Figure S1_Ferrareze et al 2020.pptx

### Slide 1
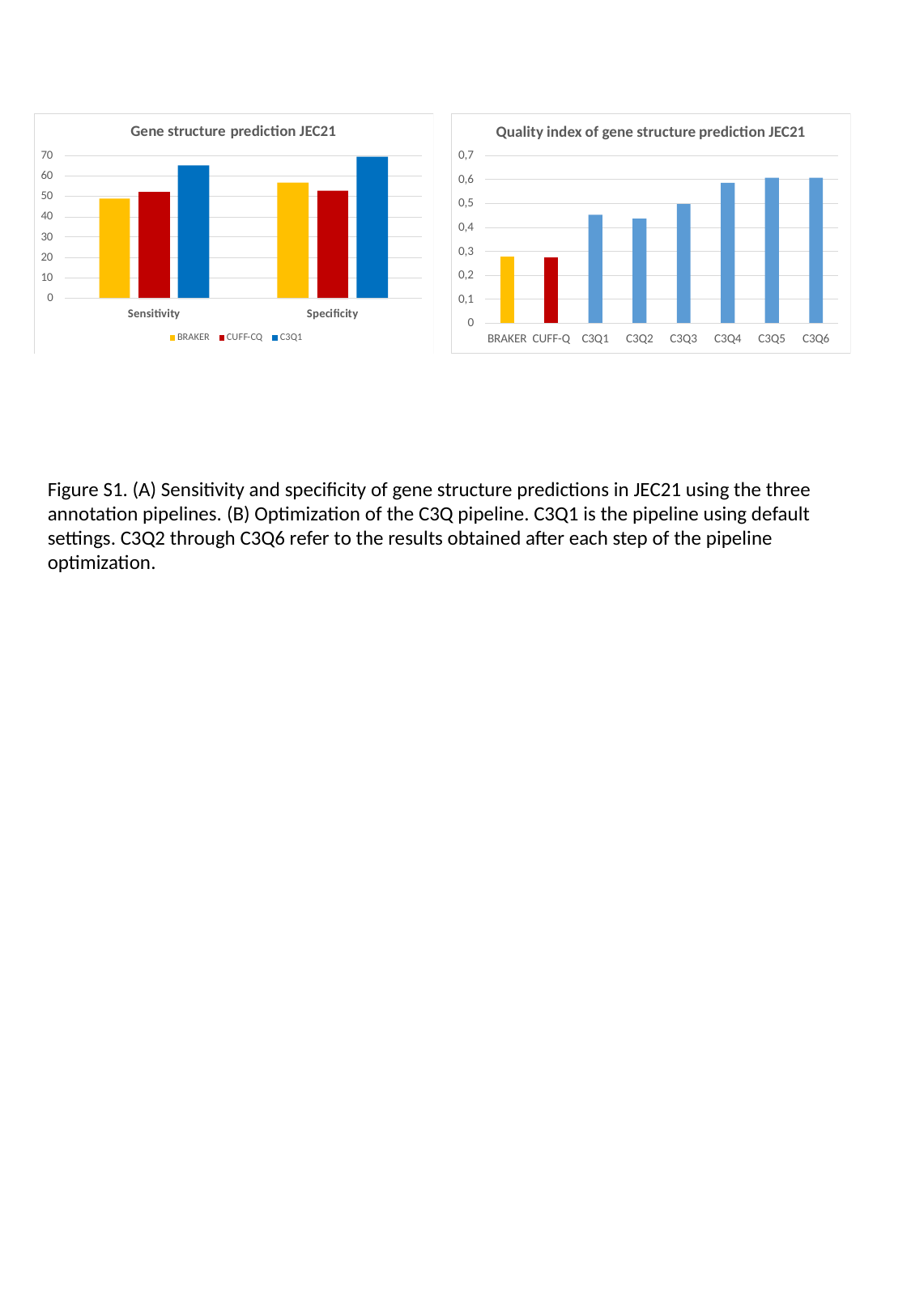

Figure S1. (A) Sensitivity and specificity of gene structure predictions in JEC21 using the three annotation pipelines. (B) Optimization of the C3Q pipeline. C3Q1 is the pipeline using default settings. C3Q2 through C3Q6 refer to the results obtained after each step of the pipeline optimization.
