## Supplementary material for "Application of an Optimized Annotation Pipeline to the *Cryptococcus Deuterogattii* Genome Reveals Dynamic Primary Metabolic Gene Clusters and Genomic Impact of RNAi Loss": Figure S2_Ferrareze et al. 2020.pptx

### Slide 1
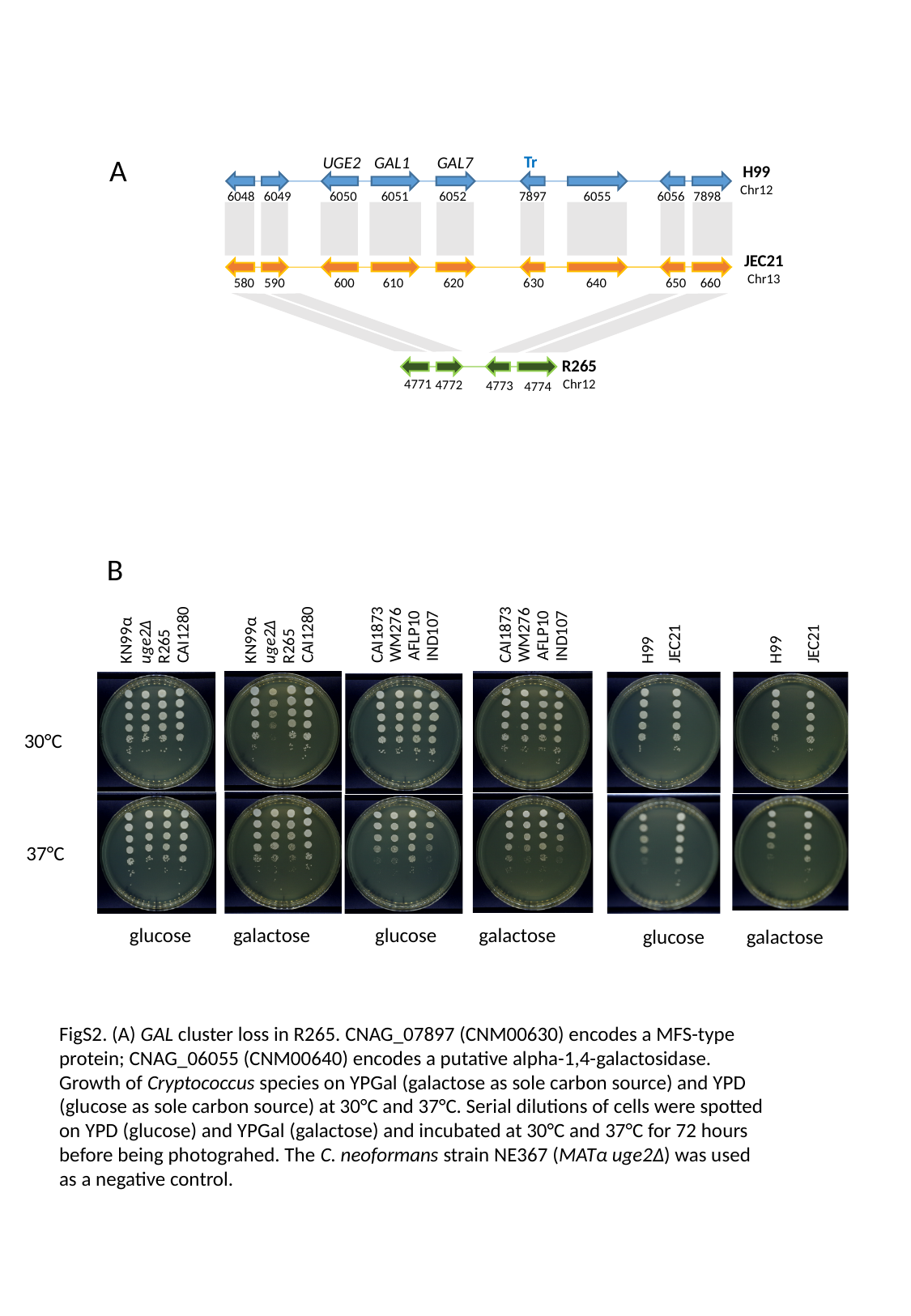

A
Tr
UGE2
GAL1
GAL7
H99
Chr12
6048
6049
6050
6051
6052
7897
6055
6056
7898
JEC21
Chr13
580
590
600
610
620
630
640
650
660
R265
Chr12
4771
4772
4773
4774
B
CAI1280
KN99α
uge2Δ
R265
CAI1280
KN99α
uge2Δ
R265
CAI1873
WM276
AFLP10
IND107
CAI1873
WM276
AFLP10
IND107
JEC21
JEC21
H99
H99
30°C
37°C
glucose
galactose
glucose
galactose
glucose
galactose
FigS2. (A) GAL cluster loss in R265. CNAG_07897 (CNM00630) encodes a MFS-type protein; CNAG_06055 (CNM00640) encodes a putative alpha-1,4-galactosidase. Growth of Cryptococcus species on YPGal (galactose as sole carbon source) and YPD (glucose as sole carbon source) at 30°C and 37°C. Serial dilutions of cells were spotted on YPD (glucose) and YPGal (galactose) and incubated at 30°C and 37°C for 72 hours before being photograhed. The C. neoformans strain NE367 (MATα uge2Δ) was used as a negative control.
