## Supplementary material for "Application of an Optimized Annotation Pipeline to the *Cryptococcus Deuterogattii* Genome Reveals Dynamic Primary Metabolic Gene Clusters and Genomic Impact of RNAi Loss": Figure S3_Ferrareze et al. 2020.pptx

### Slide 1
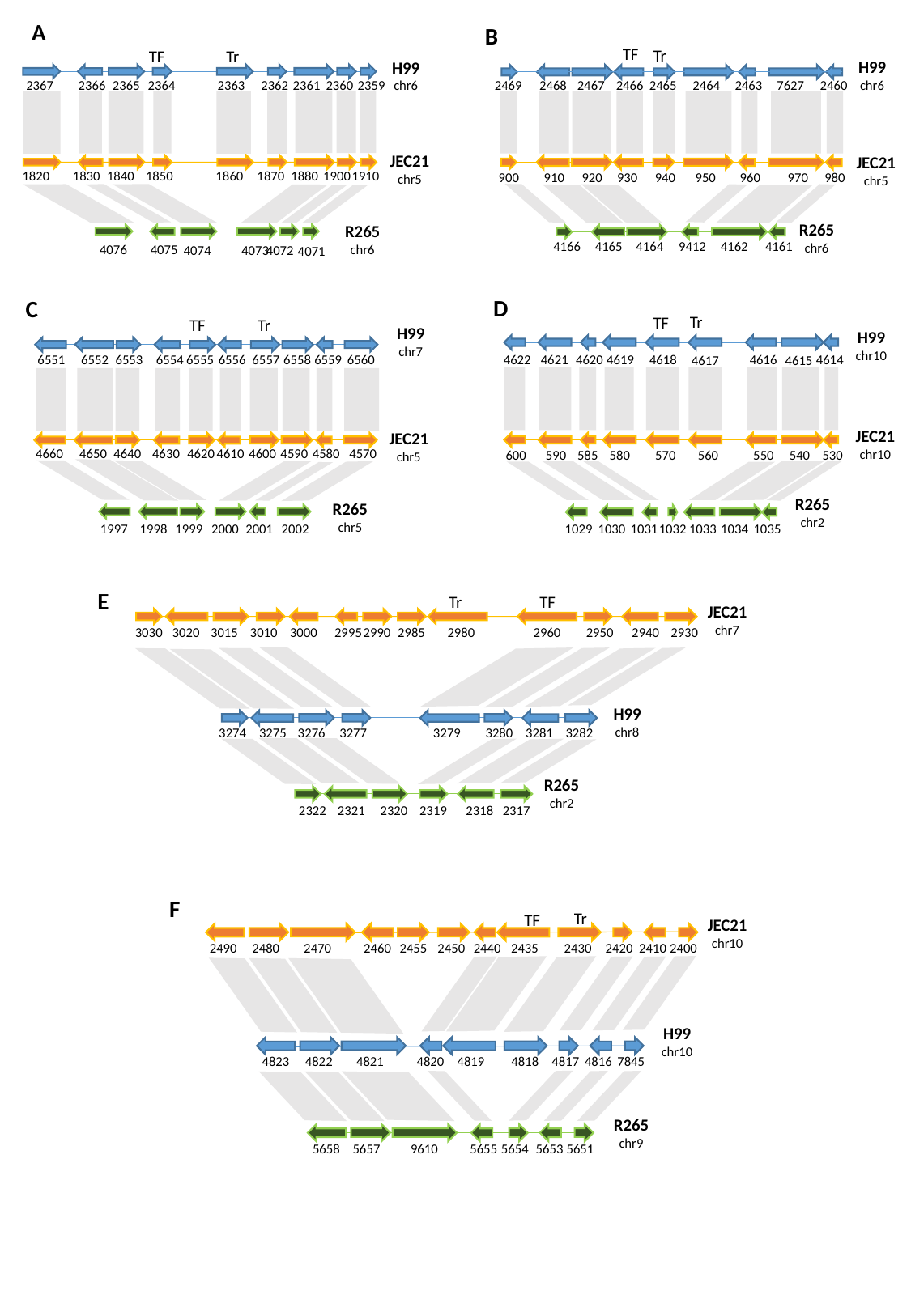

A
B
TF
Tr
Tr
TF
H99
chr6
2469
2468
2467
2466
2465
2464
2463
7627
2460
JEC21
chr5
900
910
920
930
940
950
960
970
980
R265
chr6
4166
4165
4164
9412
4162
4161
H99
chr6
2367
2366
2365
2364
2363
2362
2361
2360
2359
JEC21
chr5
1820
1830
1840
1850
1860
1870
1880
1900
1910
R265
chr6
4076
4075
4073
4072
4074
4071
D
C
Tr
TF
Tr
TF
H99
chr7
6551
6552
6553
6554
6555
6556
6557
6558
6559
6560
JEC21
chr5
4620
4660
4650
4640
4630
4610
4600
4590
4580
4570
R265
chr5
1997
1998
1999
2000
2001
2002
H99
chr10
4622
4621
4616
4620
4619
4618
4614
4617
4615
600
590
585
580
570
560
550
540
530
1029
1030
1031
1032
1033
1034
1035
JEC21
chr10
R265
chr2
E
TF
Tr
JEC21
chr7
3030
3020
3015
3010
3000
2995
2990
2985
2980
2960
2950
2930
2940
3274
3275
3276
3277
3279
3280
3281
3282
2322
2321
2320
2319
2318
2317
H99
chr8
R265
chr2
F
Tr
TF
JEC21
chr10
2490
2480
2470
2460
2455
2450
2440
2435
2430
2420
2410
2400
4823
4822
4821
4820
4819
4818
4817
4816
7845
5658
5657
9610
5655
5654
5653
5651
H99
chr10
R265
chr9

### Slide 2
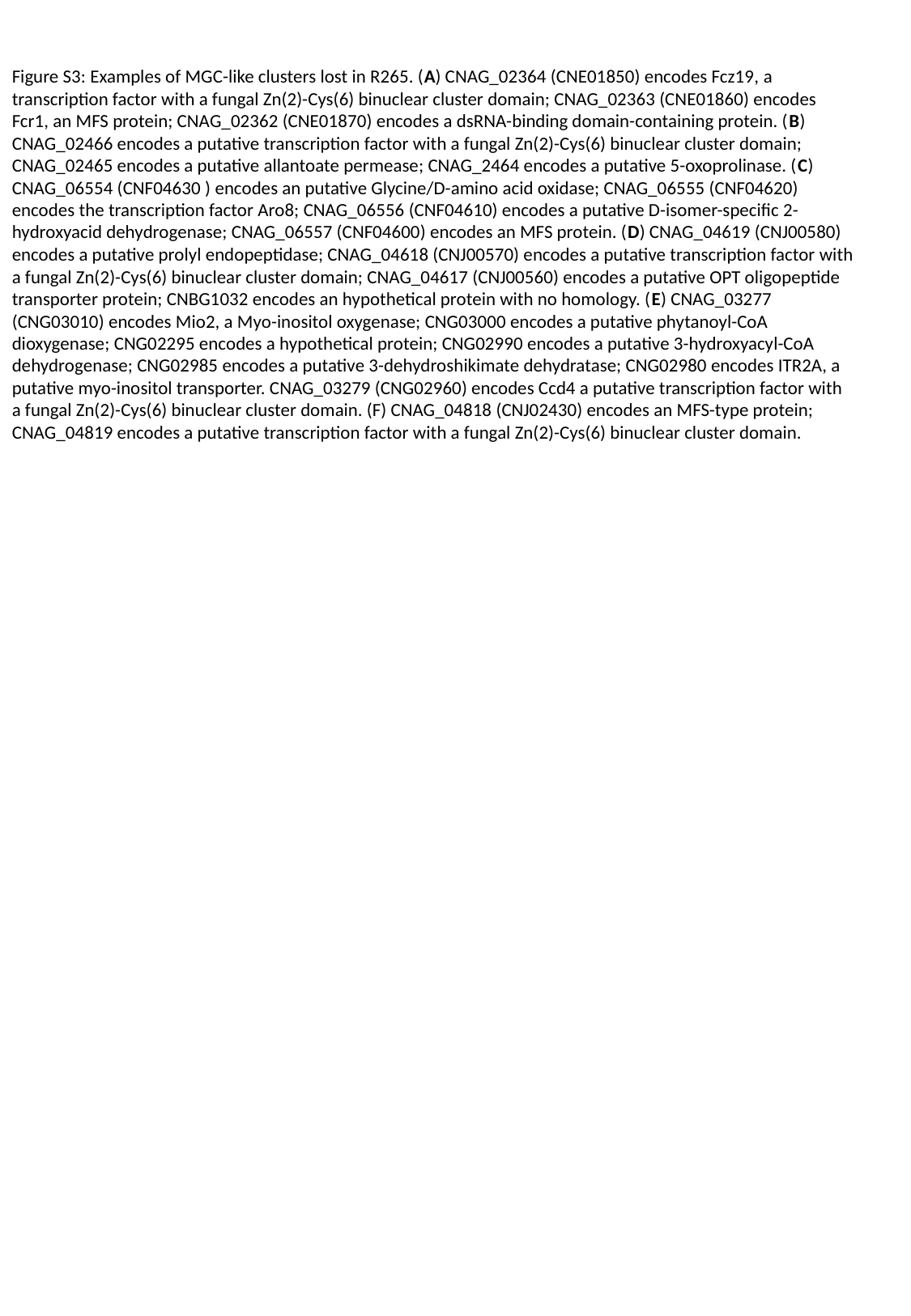

Figure S3: Examples of MGC-like clusters lost in R265. (A) CNAG_02364 (CNE01850) encodes Fcz19, a transcription factor with a fungal Zn(2)-Cys(6) binuclear cluster domain; CNAG_02363 (CNE01860) encodes Fcr1, an MFS protein; CNAG_02362 (CNE01870) encodes a dsRNA-binding domain-containing protein. (B) CNAG_02466 encodes a putative transcription factor with a fungal Zn(2)-Cys(6) binuclear cluster domain; CNAG_02465 encodes a putative allantoate permease; CNAG_2464 encodes a putative 5-oxoprolinase. (C) CNAG_06554 (CNF04630 ) encodes an putative Glycine/D-amino acid oxidase; CNAG_06555 (CNF04620) encodes the transcription factor Aro8; CNAG_06556 (CNF04610) encodes a putative D-isomer-specific 2-hydroxyacid dehydrogenase; CNAG_06557 (CNF04600) encodes an MFS protein. (D) CNAG_04619 (CNJ00580) encodes a putative prolyl endopeptidase; CNAG_04618 (CNJ00570) encodes a putative transcription factor with a fungal Zn(2)-Cys(6) binuclear cluster domain; CNAG_04617 (CNJ00560) encodes a putative OPT oligopeptide transporter protein; CNBG1032 encodes an hypothetical protein with no homology. (E) CNAG_03277 (CNG03010) encodes Mio2, a Myo-inositol oxygenase; CNG03000 encodes a putative phytanoyl-CoA dioxygenase; CNG02295 encodes a hypothetical protein; CNG02990 encodes a putative 3-hydroxyacyl-CoA dehydrogenase; CNG02985 encodes a putative 3-dehydroshikimate dehydratase; CNG02980 encodes ITR2A, a putative myo-inositol transporter. CNAG_03279 (CNG02960) encodes Ccd4 a putative transcription factor with a fungal Zn(2)-Cys(6) binuclear cluster domain. (F) CNAG_04818 (CNJ02430) encodes an MFS-type protein; CNAG_04819 encodes a putative transcription factor with a fungal Zn(2)-Cys(6) binuclear cluster domain.
