## Supplementary material for "Application of an Optimized Annotation Pipeline to the *Cryptococcus Deuterogattii* Genome Reveals Dynamic Primary Metabolic Gene Clusters and Genomic Impact of RNAi Loss": Figure S4_Ferrareze et al. 2020.pptx

### Slide 1
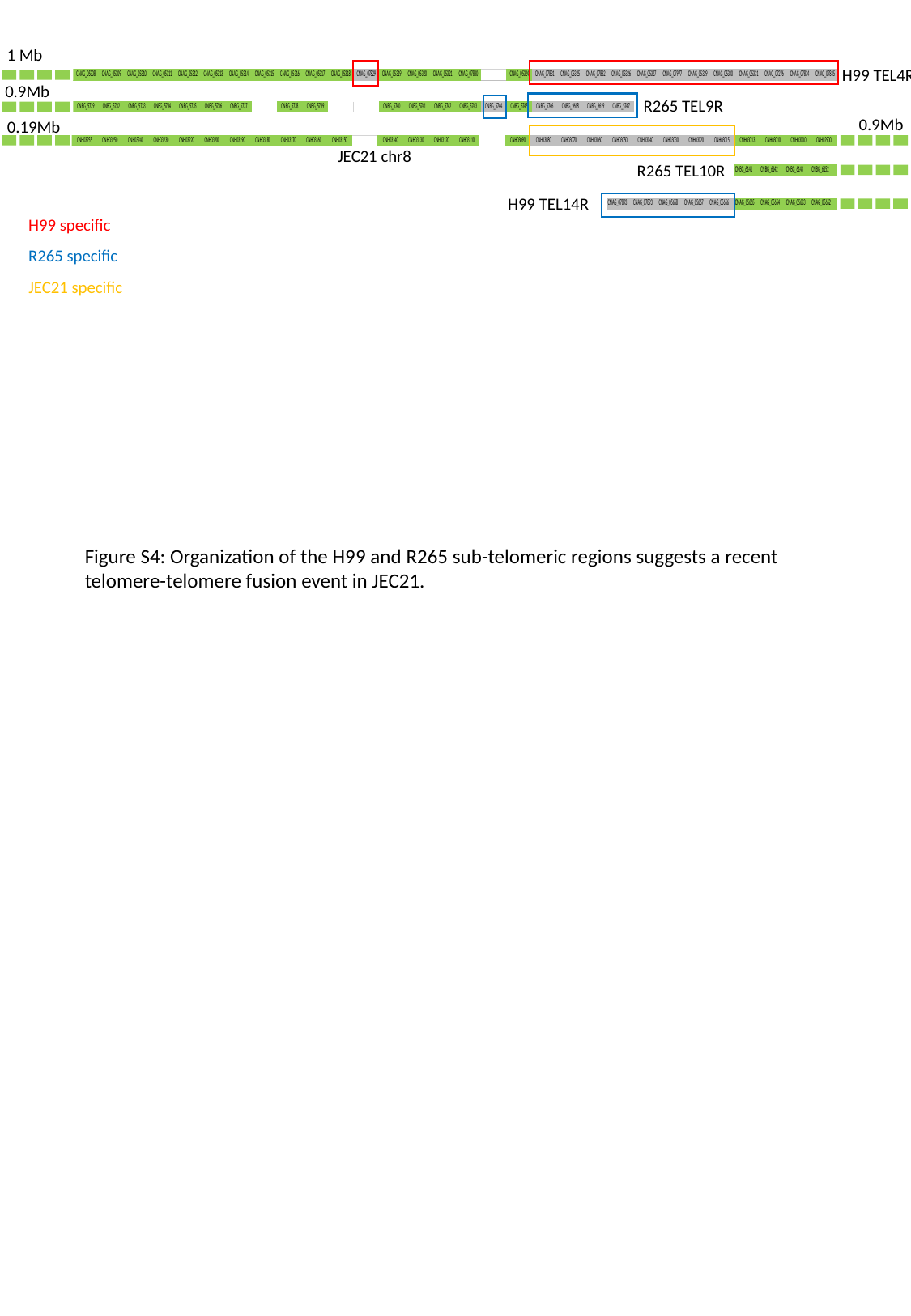

1 Mb
H99 TEL4R
0.9Mb
R265 TEL9R
0.9Mb
0.19Mb
JEC21 chr8
R265 TEL10R
H99 TEL14R
H99 specific
R265 specific
JEC21 specific
Figure S4: Organization of the H99 and R265 sub-telomeric regions suggests a recent telomere-telomere fusion event in JEC21.
